# Newly discovered base barrier cells provide compartmentalization of choroid plexus, brain and CSF

**DOI:** 10.1101/2024.07.08.601696

**Authors:** Daan Verhaege, Clint De Nolf, Jonas Castelein, Wouter Claeys, Elien Van Wonterghem, Griet Van Imschoot, Pieter Dujardin, Ward De Spiegelaere, Esther Hoste, Fleur Boone, Hart G. W. Lidov, Dani Neil, Julia Derk, Anna Kremer, Evelien Van Hamme, Peter Borghgraef, Saskia Lippens, Maria K Lehtinen, Julie Siegenthaler, Lien Van Hoecke, Roosmarijn E. Vandenbroucke

## Abstract

The choroid plexus (ChP) is a highly understudied structure of the central nervous system (CNS). The structure hangs in the brain ventricles, is composed of an epithelial cell layer, which produces the cerebrospinal fluid (CSF) and forms the blood-CSF barrier. It encapsulates a stromal mix of fenestrated capillaries, fibroblasts and a broad range of immune cells. Here, we report that the ChP base region harbors unique fibroblasts that cluster together, are connected by tight junctions and seal the ChP stroma from brain and CSF, thereby forming ChP base barrier cells (ChP BBCs). ChP BBCs are derived from meningeal mesenchymal precursors, arrive early during embryonic development, are maintained throughout life and are conserved across species. Moreover, we provide transcriptional profiles and key markers to label ChP BBCs and observe a striking transcriptional similarity with meningeal arachnoid barrier cells (ABCs). Finally, we provide evidence that this fibroblast cluster functions as a barrier to control communication between CSF and the ChP stroma and between the latter and the brain parenchyma. Moreover, loss of barrier function was observed during an inflammatory insult. Altogether, we have identified a novel barrier that provides functional compartmentalization of ChP, brain and CSF.

**GRAPHICAL ABSTRACT:** Newly discovered base barrier cells provide compartmentalization of choroid plexus, brain and CSF
The choroid plexus (ChP) hangs in the brain ventricles and is composed of an epithelial cell layer which produces the cerebrospinal fluid (CSF) and forms the blood-CSF barrier. The ChP epithelial cells are continuous with the ependymal cells lining the ventricle wall. At this base region, we identified and characterized a novel subtype of fibroblasts coined the ChP base barrier cells (BBCs). ChP BBCs express tight junctions (TJs), cluster together and seal the ChP stroma from CSF and brain parenchyma. The subarachnoid space (SAS) CSF penetrates deep into choroid plexus invaginations where it is halted by ChP BBCs.
Abbreviations: E9-16.5 (embryonic day 9-16.5); P1-4 (postnatal day 1-4).

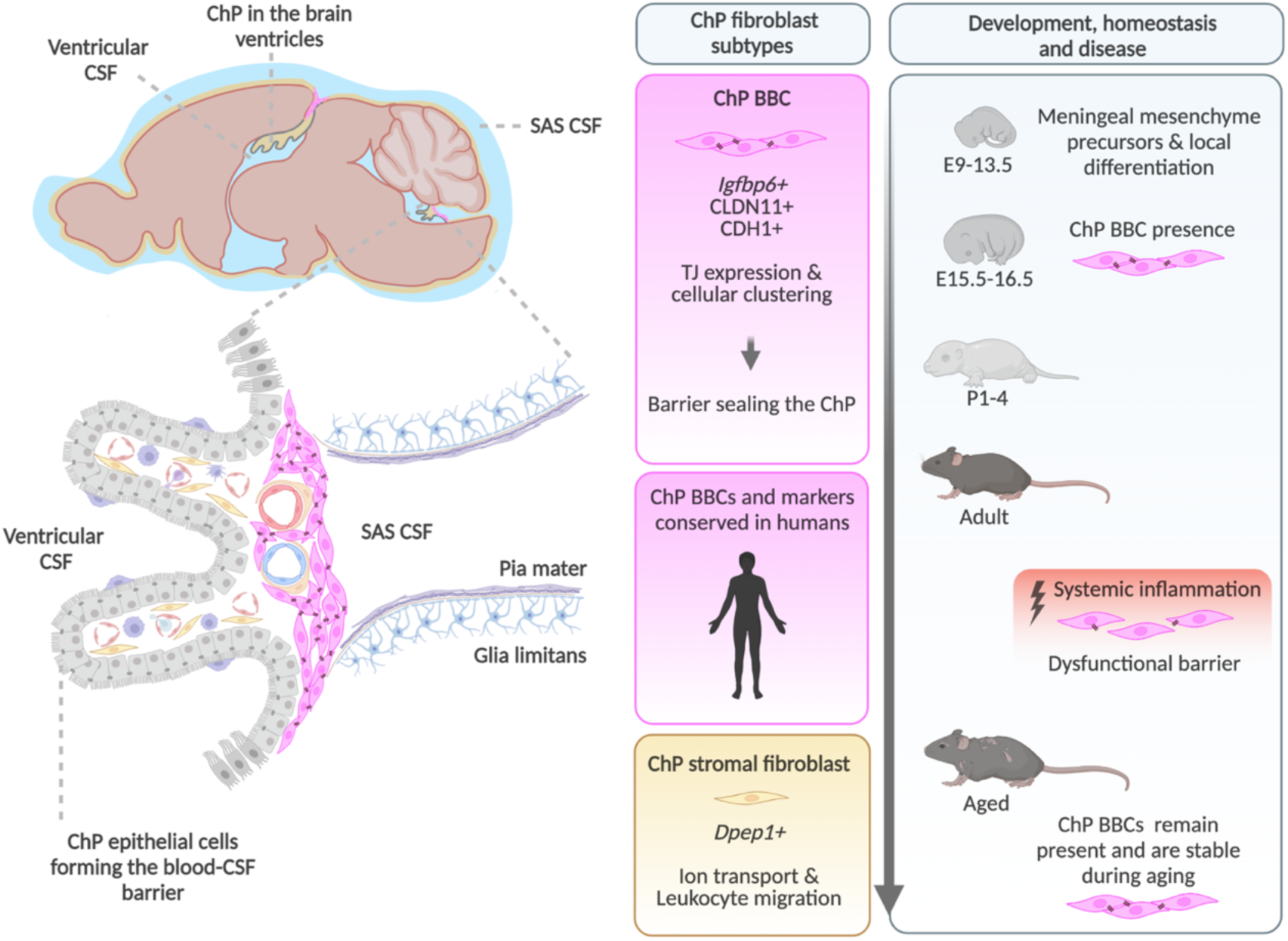

## INTRODUCTION

Tight barriers separate the central nervous system (CNS) from the periphery, assuring a balanced and well controlled micro-environment in the CNS, and providing protection against external insults such as toxins and infectious agents. The most studied brain barrier, the blood-brain-barrier (BBB), is located at the extensive brain vascular network^1^. It is composed of endothelial cells, pericytes, astrocytes and neurons that function as one neurogliovascular unit. The brain endothelial cells express tight junctions and unique transporters with limited vesicular transport and lack of adhesion molecules to ensure a tight BBB^2^. Although the BBB has been the main focus of study for years, additional but overlooked brain barriers have major roles in maintaining brain homeostasis. These barriers are located at the interface between the peripheral circulation and cerebrospinal fluid (CSF), such as the outer/dural layer of the meninges and the choroid plexus (ChP) in the brain ventricles, both contributing to the so-called blood-CSF barrier.

The meninges that enclose the brain are specified into three layers: the pia mater covering the surface of the brain parenchyma, the arachnoid mater traversing the CSF-filled subarachnoid space (SAS) and the outer dura mater consisting of fibroblasts and collagens that connect the meninges to the skull. This dural layer is further characterized by the presence of fenestrated blood vessels, facilitating the entry of blood content and immune cells. However, communication with the underlying CSF is limited by the presence of the arachnoid barrier cells (ABCs). ABCs are meningeal cells which are interconnected through tight junctions and display many epithelial-like features^3^. An additional blood-CSF interface is located at the ChP, a small organ floating in each of the four brain ventricles and mainly known for the production and detoxification of the CSF^4^. The ChP contains a vast network of fenestrated capillaries, however, free movement of molecules and cells into the CSF is prevented by tight junctions in between the ChP epithelial cells thereby ensuring regulated transport between ChP stroma and the CSF.

One striking anatomical feature of the ChP that has been largely overlooked by the field, is that the ChP stromal compartment is a direct continuation of the meninges. Studies in the developing avian chick suggest that meningeal tissue moves into the ChP stromal compartment^5^ and might give rise to the fibroblast populations of the ChP, although cellular and molecular evidence is lacking. In mice, the developing E9-E10 CNS is first enveloped by a layer of mesenchymal cells derived from either the somatic and cephalic mesoderm (for the midbrain, hindbrain and spinal cord) or the neural crest (for the forebrain)^6–9^. This initial layer of mesenchymal cells will give rise to the mesenchymal cells of the dura, arachnoid and pia mater layer fibroblasts by E14, at which point meningeal fibroblasts express unique, layer specific markers^10^. It is unknown if this early mesenchymal cell layer also contributes to the fibroblasts in the ChP, which are present in fetal brain^11^. In fact, other than a supposed role in producing the extracellular matrix^4,12^, remarkably little is known about the ChP fibroblast origin, differentiation, heterogeneity and function during health and disease. Interestingly, a recent report highlighted the physical connection of the ChP and meningeal layers, showing that the SAS penetrates deep into the ChP invaginations^13^. Although we are starting to unravel the ChP structural anatomy, the cellular composition of this ChP-meningeal continuity is still unknown. It remains to be seen if and to what extent these different structures, namely the ChP, brain, meningeal layers and CSF, are interconnected with or separated from each other.

Here, we report that the ChP contains a heterogeneous mix of fibroblasts residing in either the ChP stromal site (ChP stromal fibroblasts) or base site where the ChP is attached to the brain and the ChP-meninges connection is located (ChP base fibroblasts). These ChP base fibroblasts arrive early during embryonic development and are derived from the meningeal mesenchyme. They are maintained throughout life and are stable during aging. At this unique interface, we observe clustering of ChP base fibroblasts which are inter-connected via the presence of tight junctions. We provide evidence that this fibroblast cluster provides functional compartmentalization of the ChP, brain and CSF. We show that this fibroblast cluster functions as a barrier that controls communication between CSF and ChP stroma, and the brain and ChP stroma. Additionally, we show these barrier properties are affected during an inflammatory insult. Finally, we show cross-species conservation of ChP base fibroblasts in humans. Taken together, our findings provide exciting evidence for a novel fibroblast subtype at the ChP base, which we coin the “ChP base barrier cells” (ChP BBCs).

## RESULTS AND DISCUSSION

### scRNA-Seq reveals a novel fibroblast subtype at the choroid plexus base

To investigate the cellular composition and transcriptional profile of the ChP, we performed scRNA-Seq analyses on lateral and fourth ventricle ChP tissues of adult mice, resulting in 11.838 sequenced cells after processing (Fig. 1a). Using the expression of previously described cell-type specific markers^11,14–16^ we could identify epithelial cells, endothelial cells, pericyte-like cells, macrophages and a mix of immune cells. Notably, unbiased clustering in populations showed the presence of two distinct clusters of fibroblasts, which we refer to as ChP Type I and II fibroblasts (Fig. 1b). In a primary attempt to unravel both subtypes and examine how they relate to other fibroblasts in the CNS, we compared their transcriptome to that of non-neuronal and non-immune CNS cells by merging our adult ChP cells dataset with publicly available scRNA-Seq datasets of the developing meninges^10^, adult vascular cells^15,17^ and adult ependymal cells^17,18^ (Fig. 1c and 1d). From this merged dataset, we selected and further subclustered the CNS fibroblasts (Fig. 1e) and found that ChP Type I fibroblasts are not overlapping with other subtypes. This implies unique roles while also being distinct from ChP Type II fibroblasts. Interestingly, ChP type II fibroblasts cluster closely together with meningeal dural fibroblasts and arachnoid barrier cells (ABCs), suggesting that these fibroblasts may be akin to specific meningeal fibroblast subtypes.

**Figure 1.**
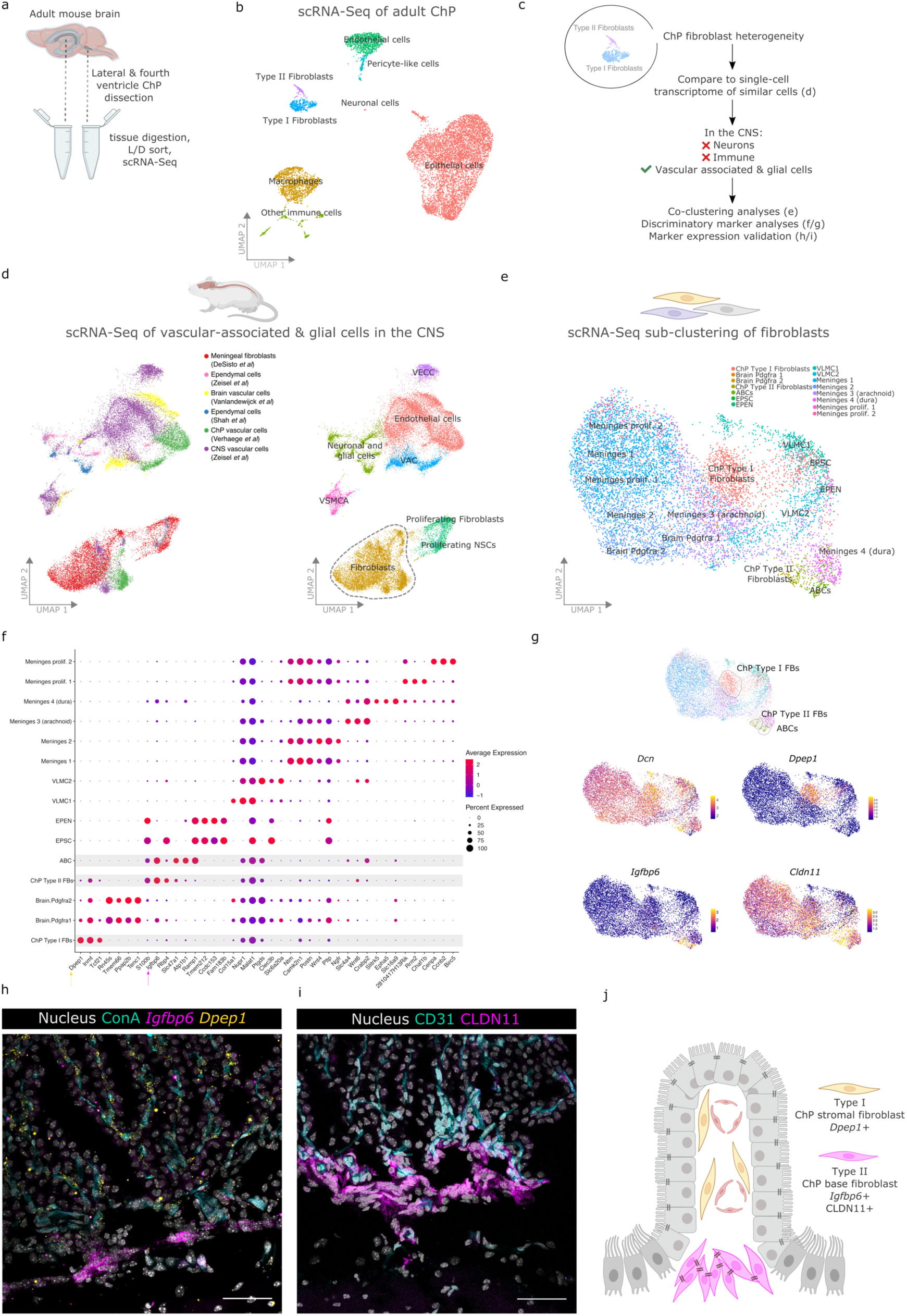
scRNA-Seq identifies a novel fibroblast subtype at the ChP base. (a) Experimental setup of lateral and fourth ventricle ChP sampling of 7-week-old mice and scRNA-Seq processing. Pooled ChP samples from N=8 mice. (b) UMAP feature plots showing the cell types detected in scRNA-Seq data of adult mouse lateral and fourth ventricle ChP (11.087 cells in total). (c) Schematic overview of selected publicly available scRNA-Seq datasets, processing and analyses. (d) UMAP feature plots showing vascular-associated and glial cells of the CNS in a merged scRNA-Seq dataset. (e) UMAP feature plots showing sub-clustering of CNS fibroblasts. (f) Dot plot representation of top 3 differential markers per CNS fibroblast subtype. (g) UMAP feature plots showing expression of Dcn, Dpep1, Igfbp6 and Cldn11 by CNS fibroblast subtypes. (h) Multiplex RNA FISH showing expression of Dpep1 (yellow) and Igfbp6 (magenta) in fourth ventricle ChP-containing brain sections obtained after ConA-FITC perfusion to label blood vessel lumen (cyan). Scale bar indicates 50 µm. Representative image for N=2 mice. (i) IF staining for CLDN11 (magenta) and CD31 (cyan) in fourth ventricle ChP-containing brain sections. Scale bar indicates 50 µm. Representative image for N=2 mice. (j) Schematic overview of location and marker expression of ChP Type I (stromal; yellow) and Type II (base; magenta) fibroblasts. ChP (choroid plexus); L/D (live/dead); CNS (central nervous system); VECC (vascular endothelial cells capillary); VSMCA (vascular smooth muscle cells arterial); VAC (Vascular associated cells/pericytes); NSC (neuronal stem cells); VLMC (vascular leptomeningeal cells); EPSC (ependymal cells spinal cord); ABC (arachnoid barrier cells)

Next, we investigated if the ChP fibroblast subtypes expressed any differential markers which allow for their discrimination from other cell (sub)types, to look into their localization in the tissue and hint at their potential functions. The differential marker analysis on CNS fibroblast subtypes (Fig. 1f and Suppl. Table 1) confirmed previously reported discriminatory markers for the various meningeal fibroblast subsets^10^, brain perivascular cells^15,17^ and ependymal cells^19,20^. In contrast to the overlap in marker expression between meningeal fibroblasts subtypes, we found striking differences between ChP Type I (expressing high levels of Dpep1, Inmt and Tcf21) and ChP Type II fibroblasts (expressing S100b and Igfbp6) (Fig. 1f and 1g). In line with the co-clustering of ChP Type II fibroblasts and ABCs, we detected shared expression of Igfbp6 and S100b while ABCs uniquely express high levels of Slc7a1, Atp1b1, and Ramp1. Interestingly, while not being a top discriminatory marker on mRNA level, we also observed high expression of Cldn11, the characteristic tight junction expressed by ABCs, in both ABCs and ChP Type II fibroblasts (Fig. 1g).

To look into the spatial distribution of ChP Type I and II fibroblasts, we validated the in vivo marker expression through multiplex RNA FISH stainings on adult mouse brain sections. We observed Dpep1 expressing cells exclusively in the stroma of the ChP of all ventricles, and more specifically closely aligning the capillary network (Fig. 1h, yellow). Excitingly, Igfbp6 expressing cells are exclusively located at the base of the ChP and close to the brain parenchyma (Fig. 1h; magenta). Due to their unique spatial localization, we refer to ChP Type I and II fibroblasts as ChP stromal and base fibroblasts, respectively. In addition to Igfbp6, we also found Cldn11 to be highly expressed by ABCs and ChP base fibroblasts and this was confirmed via immunofluorescent stainings showing CLDN11 expression at the base of the ChP (Fig. 1i and 1j; magenta).

These data reveal the existence of distinct ChP fibroblast subtypes with unique spatial localization and marker expression, namely Dpep1+ ChP stromal versus Igfbp6/CLDN11+ ChP base fibroblasts. We next investigated whether these two fibroblast subtypes have distinct functions. Interestingly, Dpep1 was recently shown to act as a physical adhesion receptor for neutrophil, monocyte and macrophage infiltration in kidney, liver and lungs^21,22^. Based on the Dpep1 expression closely aligning the ChP capillaries, it is tempting to speculate that the ChP stromal fibroblasts play an active role in the infiltration of immune cells into the ChP. Conversely, ChP base fibroblasts and ABCs share expression of Igfbp6 and Cldn11, genes responsible for insulin-like growth factor 2-induced cell proliferation/differentiation/ migration, and the formation of tight junctions, respectively^10,23^. In addition to the transcriptional similarities with ABCs, the unique localization of the ChP base fibroblasts inspired us to further explore this cell subtype.

### Choroid plexus base fibroblasts are derived from the meningeal mesenchyme, emerge early during development and are maintained throughout life

To get a better understanding of the ChP base fibroblast subtype, we studied the spatiotemporal and transcriptional profile of these cells throughout life. First, we performed CLDN11 protein stainings and analyzed mRNA in situ hybridization data of ChP base fibroblast markers in embryonic, postnatal, adult and aged mice. Clear CLDN11+ cells can be observed in the ChP base region from E16.5 onwards and these are maintained throughout life (Fig. 2a). No differences in the overall CLDN11 subcellular localization nor signal intensity was observed across ages. At E12, interpretation of CLDN11 signal is hampered by the autofluorescence of nucleated red blood cells clustering in the ChP stroma and near the base (Fig. 2a). These large and nucleated red blood cells are most likely the progenitors of the primitive erythroid lineage, which were recently described in the developing ChP^24^. To circumvent these issues, we analyzed mRNA ISH data available via the Allen Brain Atlas and observed clear Cldn11 expression in meningeal layers and the ChP base region as early as E13.5 and maintained throughout life (Fig. 2b), while Igfbp6 expression was only observed from E15.5 onwards in the same region. Together, this validates the expression of Igfbp6 and Cldn11/CLDN11 by meningeal cells reaching deep into the attachment zone of the ChP from early embryonic development onwards and being maintained throughout life (Fig. 2c).

**Figure 2.**
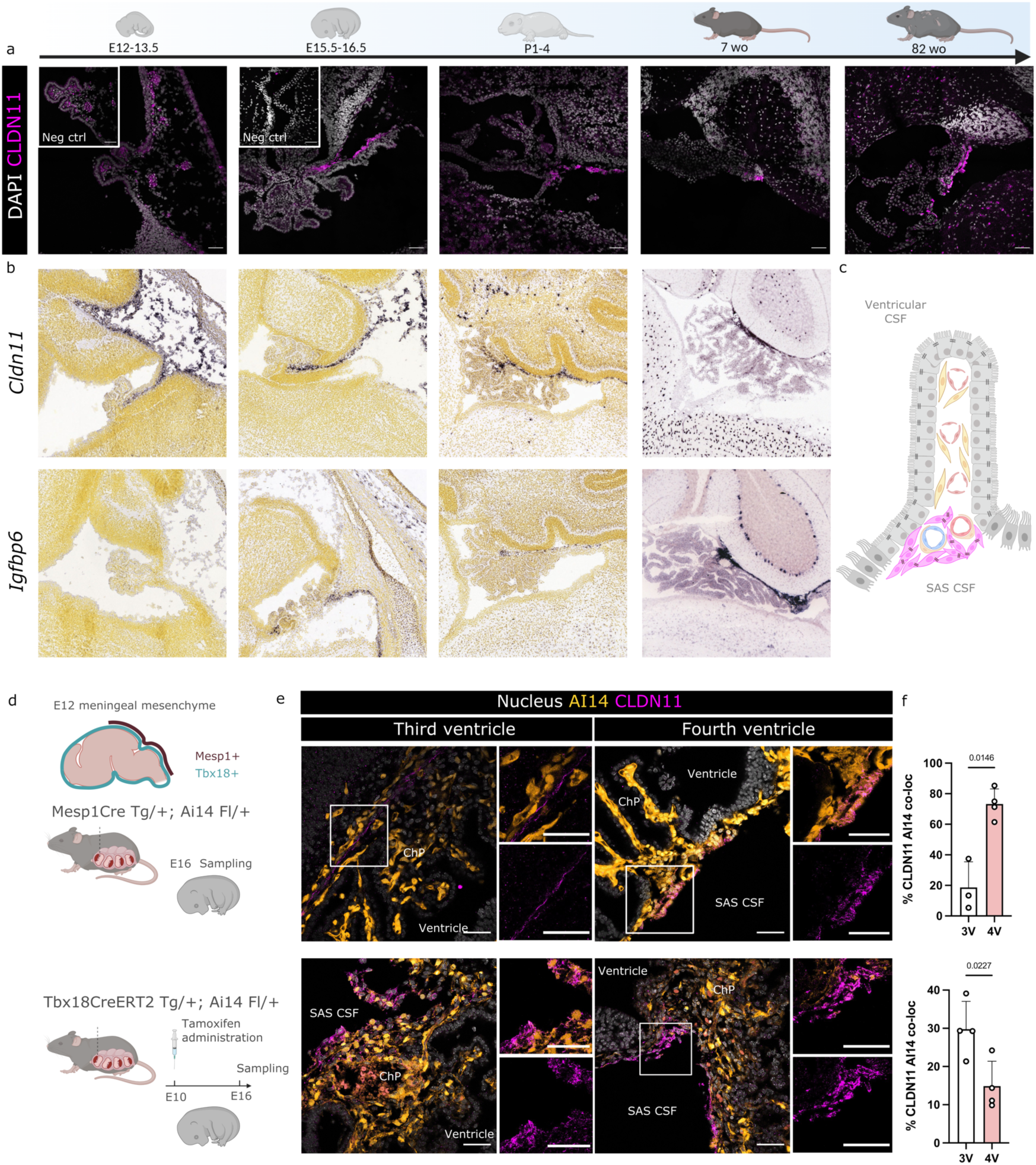
ChP base fibroblasts are present from early developmental stages and are derived from the meningeal mesenchyme. (a) IF staining for CLDN11 (magenta) in fourth ventricle ChP-containing brain sections of naive E12, E16, P1, 7-and 82-week-old mice. Scale bar indicates 50 µm. Representative image for N=2 mice per group. (b) RNA ISH showing expression of Cldn11 and Igfbp6 in fourth ventricle ChP-containing brain sections of naive E13.5, E15.5, P1 and 8-week-old mice. (adapted from the Allen Brain Atlas). (c) Schematic overview of location of ChP stromal (yellow) and base (magenta) fibroblasts, note continuity between ChP and meninges/CSF. (d) Schematic overview of transgenic lineage tracing approaches. Highlighted are targeted zones (e) IF staining for CLDN11 (magenta) in fourth and third ventricle ChP-containing brain sections of lineage tracing mice (AI14, orange) at E16. Scale bar indicates 50 µm. Representative images for N=4 mice. (f) Quantification of overlap between CLDN11 and AI14 signal in fourth and third ventricle ChP-containing brain cryosections of E16 lineage mouse brains for both tracing setups. N=4 mice per group (excluding 3V Mesp1 setup, N=3) ChP (choroid plexus); Wo (weeks old); SAS (subarachnoid space); CSF (cerebrospinal fluid); 3V (third ventricle); 4V (fourth ventricle).

The developmental origin of ChP fibroblasts is unknown. During mouse development, the brain is first surrounded by an undifferentiated mesenchymal cell population coined the primary meninx^8,9^. From E9 to E13 this layer differentiates and gives rise to the dural, leptomeningeal and pial layers, but its contribution to the ChP is unclear. Interestingly, ChP stromal fibroblasts express high levels of Tcf21, a transcription factor expressed in mesoderm and embryonic mesenchymal-derived cells (Fig. 1f)^25,26^. We also observed expression of the mesenchymal stem cell marker PDGFRα in the stroma and base of the ChP (Suppl. Fig. 1)^27^. To test the contribution of the meningeal mesenchyme as progenitors of ChP fibroblasts, we lineage traced Mesp1-Cre Tg/+ ^28^ crossed with Ai14 flox reporter mice (Fig. 2d). Mesp1-Cre is a mesoderm lineage Cre that has been previously shown to recombine in mid/hindbrain meningeal fibroblasts, but also pericytes in this brain region and all endothelial cells^29^. Interestingly, CLDN11+ cells at the 4^th^ ventricle ChP base were lineage positive, whereas those in the 3^rd^ ventricle ChP were lineage mixed (Fig. 2e and 2f), likely due to contribution of neural crest and mesoderm-derived meningeal mesenchyme in this area. These data suggest that ChP base fibroblasts in the different ventricles are derived from meningeal mesenchyme matching their nearby location during development. In order to investigate the contribution of the meningeal mesenchyme in a time-dependent manner, we used Tbx18-CreERT2 Tg/+ ^30^; Ai14 fl/+ mice (Fig. 2d). These mice display Cre expression in meningeal mesenchymal cells throughout the CNS and mural cells^10^. Using a similar approach used to show that ABCs are derived from the early meningeal mesenchyme, we injected mice with tamoxifen at E10 (Fig. 2c)^31^, a timepoint that precedes the presence or specification of both ABCs and ChP base fibroblasts. Notably, although this inducible approach resulted in more sparse labeling of CLDN11+ cells at the base of the E16 ChP as compared to the constitutively active Mesp1-Cre, there was still a clear fraction of lineage positive cells (Fig. 2e), with more cells lineage positive in the third ventricle ChP. Thus, the meningeal mesenchyme gives rise to the ChP base fibroblasts and ABCs, with their specific differentiation likely locally instructed by the niche. In line with this, Desisto et al recently performed scRNA-Seq analyses on the E14.5 developing mouse meninges and reported on vast fibroblast heterogeneity in the different layers^10^. Interestingly, they observed a transcriptional outlier population which was associated with pial layers and was found to extend into the ChP. Based on this, it is tempting to speculate that these cells will give rise to ChP base fibroblasts by E16. Noteworthy, the early emergence of ChP base fibroblasts is in stark difference to e.g. perivascular fibroblasts which only develop postnatally^32^, and begs the question about their potential roles during early development.

In addition to the spatiotemporal profile of ChP base fibroblasts emergence, development and maintenance, we looked into their transcriptomic profile throughout life. In brief, scRNA-Seq analyses were performed on lateral and fourth ventricle ChP of 7-, 22- and 82-week-old mice (18.783 and 18.222 single cells in total for 22- and 82-week-old samples, respectively), and compared to E16.5 ChP^11^ (Fig. 3a). Although several embryonic-specific fibroblast clusters are observed, the majority of fibroblasts populate the two main clusters of ChP stromal and base fibroblasts (Fig. 3b and 3c), expressing general fibroblast markers (Dcn) but again show discriminatory expression of Cldn11, Igfbp6 and Dpep1 (Fig. 3d). Notably, we observed a small cluster of most likely contaminating ABCs which cluster close to -but not with- the ChP base fibroblasts, again pointing towards shared origins and/or biological functions (Fig. 3c). In line with our spatiotemporal analysis of ChP base fibroblasts, the ChP stromal and base fibroblasts clusters are composed of cells originating from E16.5, 7-, 22- and 82-week-old mice (Fig. 3e). Next, we analyzed the expression over time of ChP base fibroblast markers (Igfbp6 and Cldn11) and ChP stromal fibroblast markers (Dpep1 and Alpl) in these clusters. At all ages, we could confirm that Igfbp6 expression is restricted to ChP base fibroblasts and Cldn11 is expressed at higher levels in ChP base compared to stromal fibroblasts (Fig. 3f). Similarly, Dpep1 expression is restricted to the ChP stromal fibroblasts and Alpl (an additional ChP stromal fibroblast marker) expression is higher in ChP stromal fibroblasts (Fig. 3f). The expression of these markers remained stable from adult stage onwards. Strikingly, no differentially expressed genes were observed when comparing ChP base fibroblasts from 22-versus 82-week-old mice. Our data show that ChP base fibroblasts originate from the meningeal mesenchyme and differentiate during developmental stage E12-E15, displaying stable expression of Igfbp6/Cldn11/CLDN11 from E15-16 onwards and throughout life (Fig. 3g).

**Figure 3.**
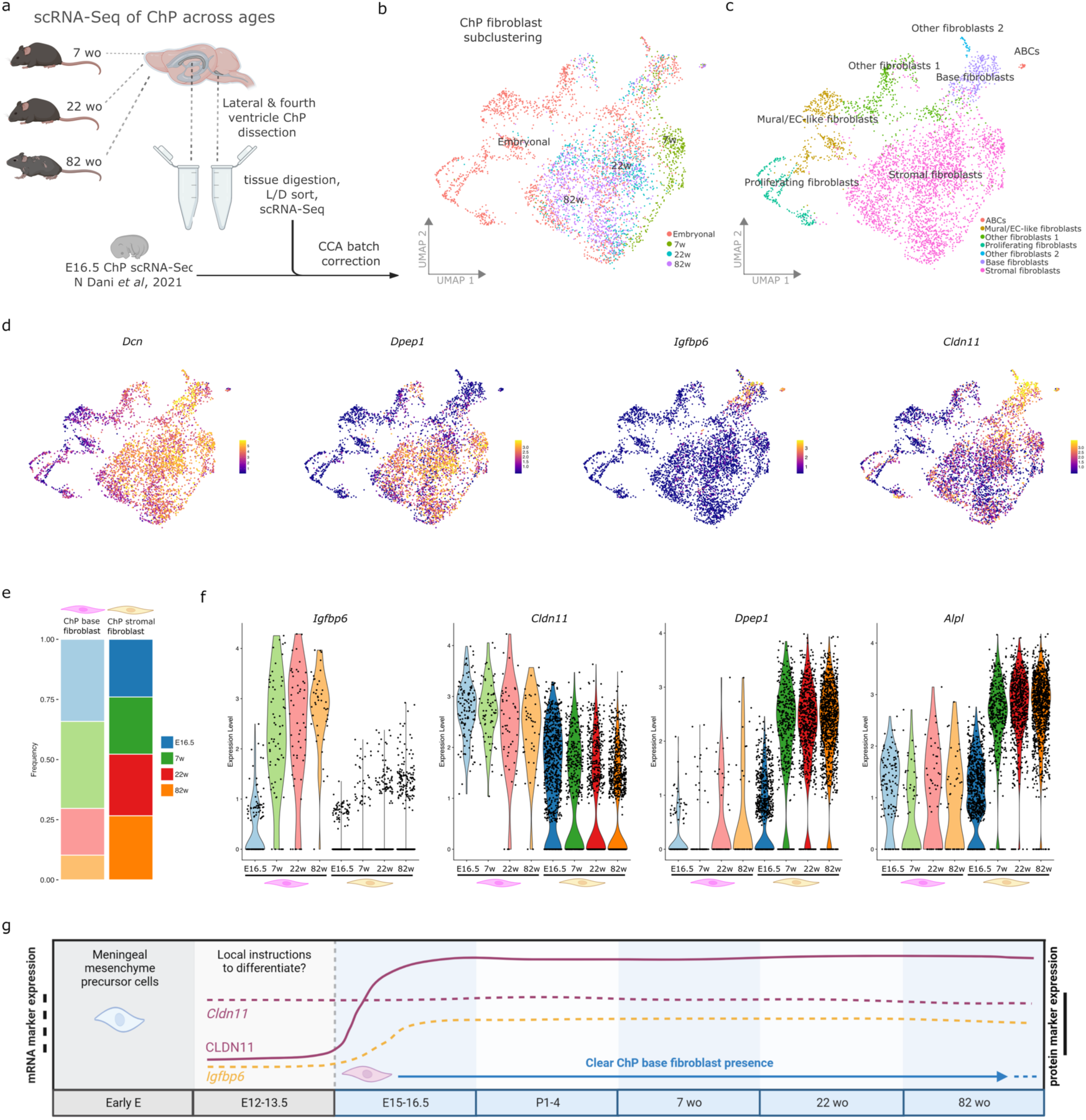
ChP base fibroblasts show stable marker expression throughout life. (a) Experimental setup of lateral and fourth ventricle ChP sampling of 7-, 22- and 82-week-old mice (N=8 pooled mice; 11.075 cells for 7-week-old samples; 18.783 cells for 22-week-old samples; 18.222 cells for 82-week-old samples), scRNA-Seq processing and combination with previously published scRNA-Seq on embryonic ChP (Dani et al, 2021). (b) UMAP feature plots showing sub-clustering of ChP fibroblasts of E16, 7-, 22- and 82-week-old mice (ages annotated). (c) UMAP feature plots showing sub-clustering of ChP fibroblasts of E16, 7-, 22- and 82-week-old mice (cell subtypes annotated). (d) UMAP feature plots showing the expression of Dcn, Dpep1, Igfbp6 and Cldn11. (e) Frequency plots showing contribution of age groups to ChP base and stromal fibroblast clusters. (f) Violin plots showing expression of Igfbp6, Cldn11, Dpep1 and Alpl in ChP stromal and base fibroblasts across ages. (g) Schematic overview summarizing the emergence, development and marker expression of ChP base fibroblasts. ChP (choroid plexus); Wo (weeks old); L/D (Live/dead); CCA (canonical correlation analysis).

### ChP base fibroblasts express tight junctions and cluster together, thereby sealing the ChP stroma from brain parenchyma and CSF

Fibroblasts throughout the body are known to produce collagens that help make up the extracellular matrix (ECM) for structural support^33^. We used our scRNA-Seq merged dataset containing CNS fibroblasts to investigate the expression of key ECM genes (Sup. Fig. 2a and 2b). This showed high expression of ECM genes by meningeal and brain perivascular cells, but only minimal expression of these genes in both ChP base and stromal fibroblasts (Sup. Fig. 2b). This was supported by Herovici-stained brain sections with overall weak and immature collagen signal in the ChP relative to control tissues (Sup. Fig. 2c). We observed no distinct collagen signal in the ChP base region. Similarly, in serial-block-face scanning EM (SBF-SEM) and focused-ion-beam (FIB) SEM (FIB-SEM) analyses of the ChP base region, we only occasionally observed thin collagen bundles (Sup. Fig. 2d). Together, these data suggest that the ChP base fibroblasts do not produce substantial amounts of ECM typical for structural support.

Next, we used the scRNA-Seq transcriptome of the ChP fibroblast subtypes to investigate their functions. Gene Ontology Enrichment Analysis (GOEA) was performed on differentially expressed genes between ChP stromal and base fibroblasts (Fig. 4a, Suppl. Table 2). GO terms associated with ChP stromal fibroblasts are 1) iron/transition metal/copper ion transport (Steap4, Heph and Cp), 2) integrin activation and leukocyte migration (Cxcl12) and 3) anchored component of the membrane with a surprising link to Dpep1 expression (Fig. 4a and 4b). Additionally, since fibroblasts utilize a broad range of integrins to coordinate their microenvironment^34^, we investigated the expression of integrins and found Itga8 to be uniquely expressed by the ChP stromal fibroblasts and associated with many of the top GO terms (Fig. 4b). This gene is linked to GO terms cell-matrix adhesion, substrate adhesion-dependent cell spreading and was recently shown to play a major role in the migration of mesenchymal cells into epithelial structures^35,36^. In conclusion, these GO analyses hint towards a situation where ChP stromal fibroblasts are surrounded by an epithelial cell layer and are closely aligning fenestrated capillaries, where they may play a role in ion transport, anchoring of membranes and leukocyte migration.

**Figure 4.**
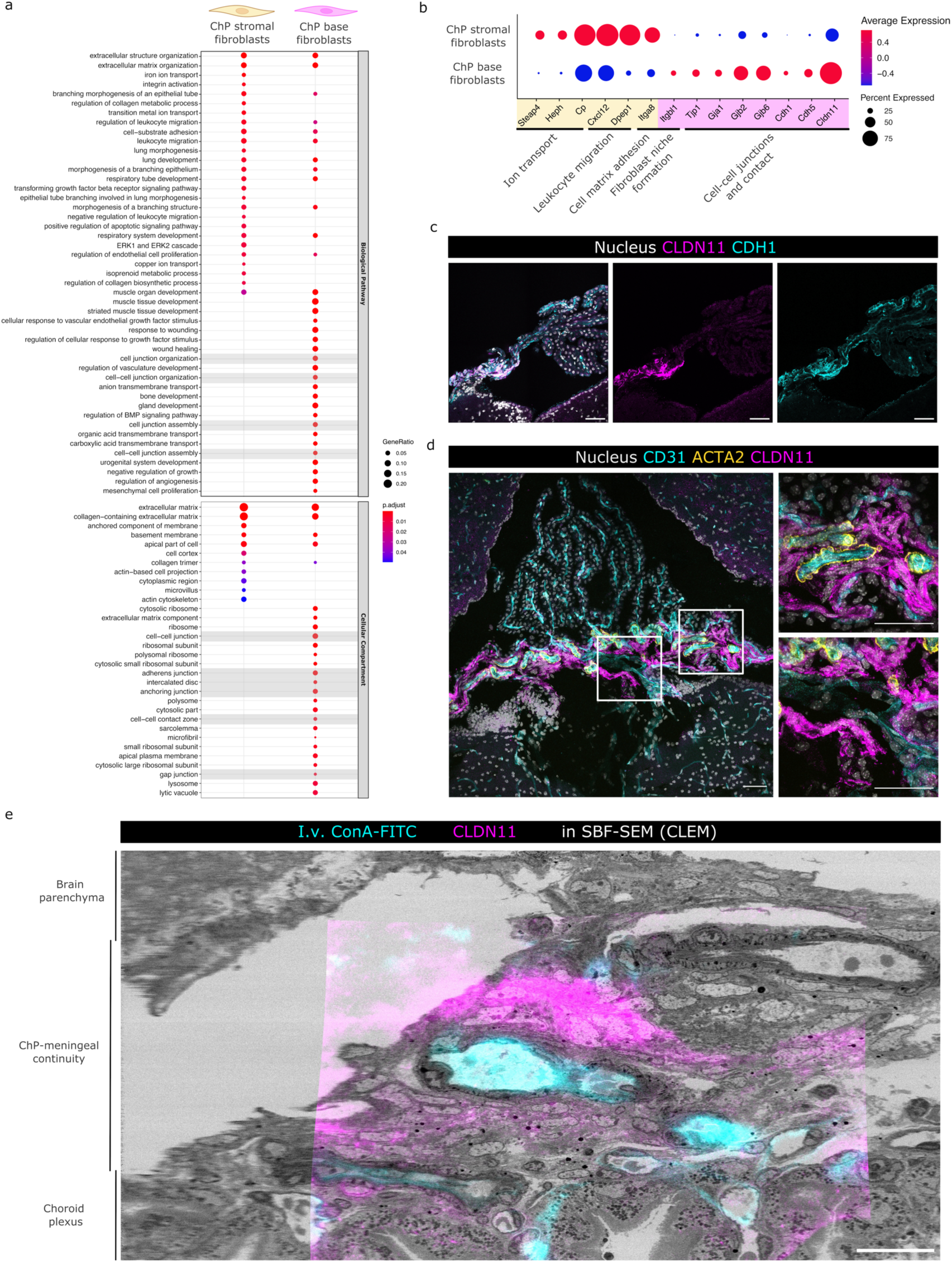
ChP base barrier cells (ChP BBCs) express tight junctions and cluster around blood vessels. (a) Gene Ontology Enrichment Analysis on Differentially Expressed genes between ChP stromal and base fibroblasts of 7-week-old mice. Highlighted in grey are GO terms unique to ChP base fibroblasts. (b) Dot plot visualization showing expression of selected discriminatory genes and their associated GO terms (c) IF staining for CLDN11 (magenta) and CDH1 (cyan) in fourth ventricle ChP-containing brain sections of 7-week-old mice. Scale bar indicates 50 µm. Representative image for N=3 mice. (d) IF staining for CLDN11 (magenta), CD31 (cyan) and ACTA2 (yellow) in fourth ventricle ChP-containing brain sections of 7-week-old mice. Scale bar indicates 50 µm. Representative image for N=4 mice. (e) IF staining for CLDN11 (magenta) on fourth ventricle ChP-containing brain vibratome sections of i.v. ConA-FITC (cyan) injected 7-week-old mice. IF microscopy was followed by CLEM showing overlap of CLDN11 IF staining signal with SBF SEM. Scale bar indicates 10 µm. Snapshot from Suppl. Movie 1. Data from single mouse. ChP (choroid plexus); SBF-SEM (serial-block-face scanning electron microscopy); CLEM (correlative light electron microscopy).

Conversely, top GO terms associated with ChP base fibroblasts are cell-cell junction organization/assembly (Cd9, Ramp2, S100a10, Gjb2, Cdh1, Cdh5), cell-cell contact zone (Tjp1, Gja1, Gjb2, Gjb6, Cdh1, Cdh5), adherens junctions, anchoring junction, gap junction and intercalated disc (Fig. 4a and 4b, highlighted in grey), hinting towards extensive cell-cell contacts made by these cells. Additionally, Cldn11 was associated with many of these pathways. Although no integrins were linked to the top 25 GO terms, the integrin Itgbl1 is uniquely expressed by the ChP base fibroblasts (Fig 4b). This gene is linked to GO terms cell-substrate adhesion, cell-matrix adhesion and was recently described to play a role in the formation of fibroblast niches^37^. To investigate this potential cell-cell contacting fibroblast niche, we analysed tight junction expression patterns in more detail. CLDN11+ CDH1+ cells are observed at the base of the ChP (Fig. 4c). Notably, we did not observe a honeycomb pattern as is often reported for tight junctions, but a rather diffuse CLDN11 expression pattern (Fig. 4d). Based on the large CLDN11+ surface at the ChP base, we hypothesized that many CLDN11+ fibroblasts cluster together closely interacting with one another. By investigating additional vasculature markers, we observed that CLDN11+ cells cluster mainly around arterioles (CD31+ ACTA2+) and to a lesser extent venules (CD31+ ACTA2-) that enter and leave the ChP tissue, respectively (Fig. 4d). In this niche, they form a dense cellular cluster interconnected through CLDN11+ CDH1+ tight junctions.

Next, high resolution visualization of the ChP base fibroblasts and their niche through electron microscopy (EM) analyses was performed using correlative light-electron microscopy (CLEM)^38^, enabling us to investigate in EM data which cells express CLDN11. CLEM visualizations confirmed the presence of CLDN11+ cells located in the region between the brain and the attached ChP tissue (Fig. 4e & Suppl. Movie 1). More specifically, these cells cluster in high numbers around arterioles and venules at the base of the ChP. Strikingly, CLDN11+ cells have a very distinct nucleus appearing large, electron sparse (pale appearance) with an outer rim of electron dense heterochromatin (Suppl. Fig. 3a and 3b). These cells could also be found using SBF-SEM analysis (Fig. 5a). Indeed, tracing these cells followed by 3D reconstruction showed clear clustering of the ChP base fibroblasts that form a plug that seals the stroma of the ChP (Fig. 5a & Suppl. Movie 2). Finally, FIB SEM analysis allowed the visualization of the cell-cell contacts between ChP base fibroblasts^39^. This revealed their cell membranes are largely in contact with one another and are connected through electron-dense cell-cell contact zones reminiscent of tight-junction organizations previously described for arachnoid barrier cells^40^ (Fig. 5b). Again, the ChP base fibroblast localization in between arterioles and venules suggests these cells might play a role in sealing the route from ChP stroma to the brain parenchyma. Simultaneously, they also prevent contact between ChP stroma and the CSF in the SAS which surrounds the brain but is also present in the ChP invagination (Fig. 5b). In line with this, we observed ramified, elongated cells at the CSF/meninges side, which were reminiscent of leptomeningeal cells traversing the SAS (Fig. 5b)^41^. Additionally, through intracerebroventricular (i.c.v.) injection into the lateral ventricle of 3 kDa biotinylated dextran, we observed that these dextrans could bind the apical surface of the ChP epithelium, but also traverse the ventricular system to reach the SAS (Figure 5c). At the SAS CSF side of the ChP base fibroblasts we observed dextran signal just next to the CLDN11+ base fibroblasts highlighting their close interactions with CSF on one side and the ChP stroma on the other side.

**Figure 5.**
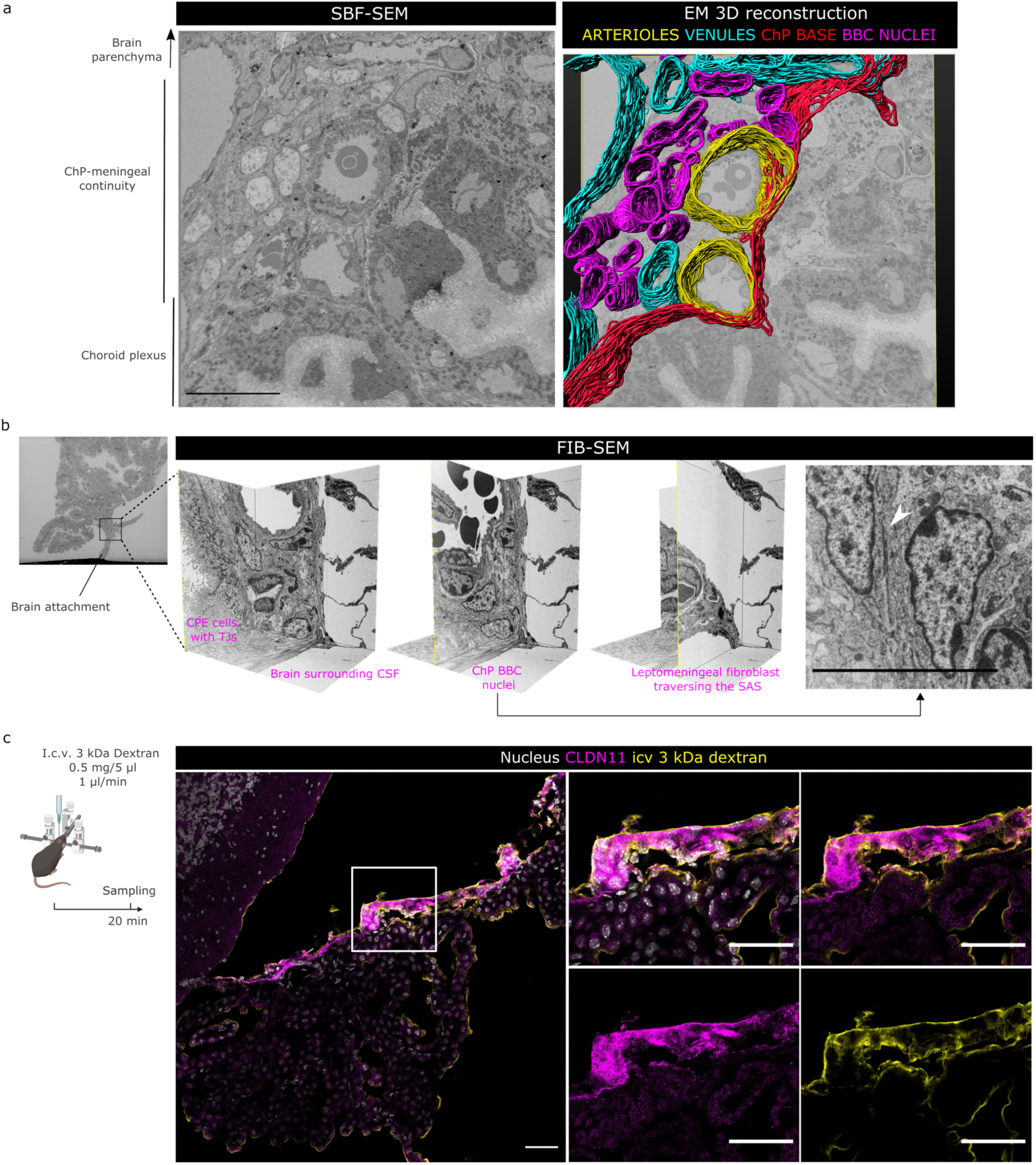
ChP base barrier cells (BBCs) seal the ChP stroma from brain and CSF. (a) SBF-SEM analyses on fourth ventricle ChP-containing brain vibratome sections (single z-plane shown, left) and 3D volume reconstruction of arterioles (yellow), venules (cyan), ChP base (red) and ChP BBC nuclei (magenta). Note plug-like BBC cluster that seals route from/towards the ChP stroma. Scale bar indicates 10 µm. Snapshot of Suppl. Movie 2. Data from single mouse. (b) FIB-SEM analyses on fourth ventricle ChP-containing brain vibratome sections (shown left). Three z-planes shown from 3D imaged block. Right: single z-plane of ChP base region showing two ChP BBCs connected through electron-dense cell-cell contact zones (white arrowhead). Scale bar indicates 10 µm. Data from single mouse. (c) IF staining for CLDN11 (magenta) and streptavidin (yellow) in fourth ventricle ChP-containing brain sections of 8-wee-old mice injected i.c.v. with 3 kDa biotinylated dextran. Scale bar indicates 50 µm. Representative image for N=4 mice. ChP (choroid plexus); SBF-SEM (serial-block-face scanning electron microscopy); FIB-SEM (focus-ion-beam scanning electron microscopy); CSF (cerebrospinal fluid); SAS (subarachnoid space); TJs (tight junctions); CPE (ChP epithelial cells).

Based on transcriptional similarities with ABCs, which are also located at the interface of CSF and fenestrated capillaries, it is most likely that the ChP base fibroblasts perform similar functions as ABCs albeit at a different location. In addition to CSF and stromal fenestrated capillaries, ChP base fibroblasts are also found at the site where ChP epithelial cells are physically attached to brain tissue. As such, the ChP base fibroblasts form a functional compartmentalization of ChP, brain and CSF and thus ensure that the ChP is an isolated structure. Altogether, the ChP base fibroblast transcriptome, TJ expression, cellular clustering and unique anatomical location are strong indicators for barrier properties and encouraged us to propose “ChP base barrier cells” (ChP BBCs) as name for these cells. To probe these barrier properties and investigate how they cope with insult, we explored the ChP BBCs in a mouse model for systemic inflammation.

### The ChP base barrier cells show dysfunctional barrier properties upon inflammatory insult

Under homeostatic conditions, the brain barriers separate the CNS from the periphery. As an example, CLDN11 is present in very tight bi- and tri-cellular junctions of arachnoid barrier cells (ABCs) that prevent the leakage of small molecules from the fenestrated blood vessels in the dura mater across ABCs and into the CSF and towards the brain^42^. Importantly, peripheral insults such as systemic inflammation are known to affect the barrier properties of both the BBB^43^ and blood-CSF barrier^44^. In line with these reports, a strong decrease in Cldn11 expression is observed in ChP tissue of systemically inflamed mice (Fig. 6a and 6c). To probe if this reduction is linked to dysfunctional barrier properties, we visualized the in vivo leakage of 3 kDa biotinylated dextran across the ChP BBCs of naive and systemically inflamed mice (Fig. 6a, 6b and 6d). I.v. injected dextrans can freely enter the ChP stroma through fenestrations of the capillaries but are unable to leave the blood vessels in the brain parenchyma (Fig. 6e). Only faint dextran signal is visible in the region connecting the ChP to the brain parenchyma of naive mice, suggesting only little leakage of small molecules occurs during homeostasis (Fig. 6e, arrows). However, under systemic inflammatory conditions we observed patches of intense dextran signal across the ChP base region (Fig. 6e, arrows and zoom-ins) and this leakage was associated with a significant increase in CLDN11-Dextran co-localization (Fig. 6d). As such, these in vivo analyses of barrier integrity suggest that peripheral insults such as systemic inflammation can negatively affect the ChP BBCs resulting in leakage from the ChP stroma to the brain. Although a leaky barrier does not necessarily mean that immune cell transport is facilitated, it would be interesting to investigate immune cell transport across the ChP BBCs and towards the brain or into the CSF. Indeed, monocyte, neutrophil and T cell accumulation in the ChP has been described in various mouse models for inflammation and diseases of the CNS^45–47^. Although many reports suggest these immune cells breach the ChP epithelial barrier after which they are detected in the CSF, there is only little in vivo evidence of such crossing due to technical challenges of visualizing crossing events^48^. Generating transgenic mice to target the ChP base barrier cells in combination with imaging tools will be key to unravel their potential roles in CNS diseases.

**Figure 6.**
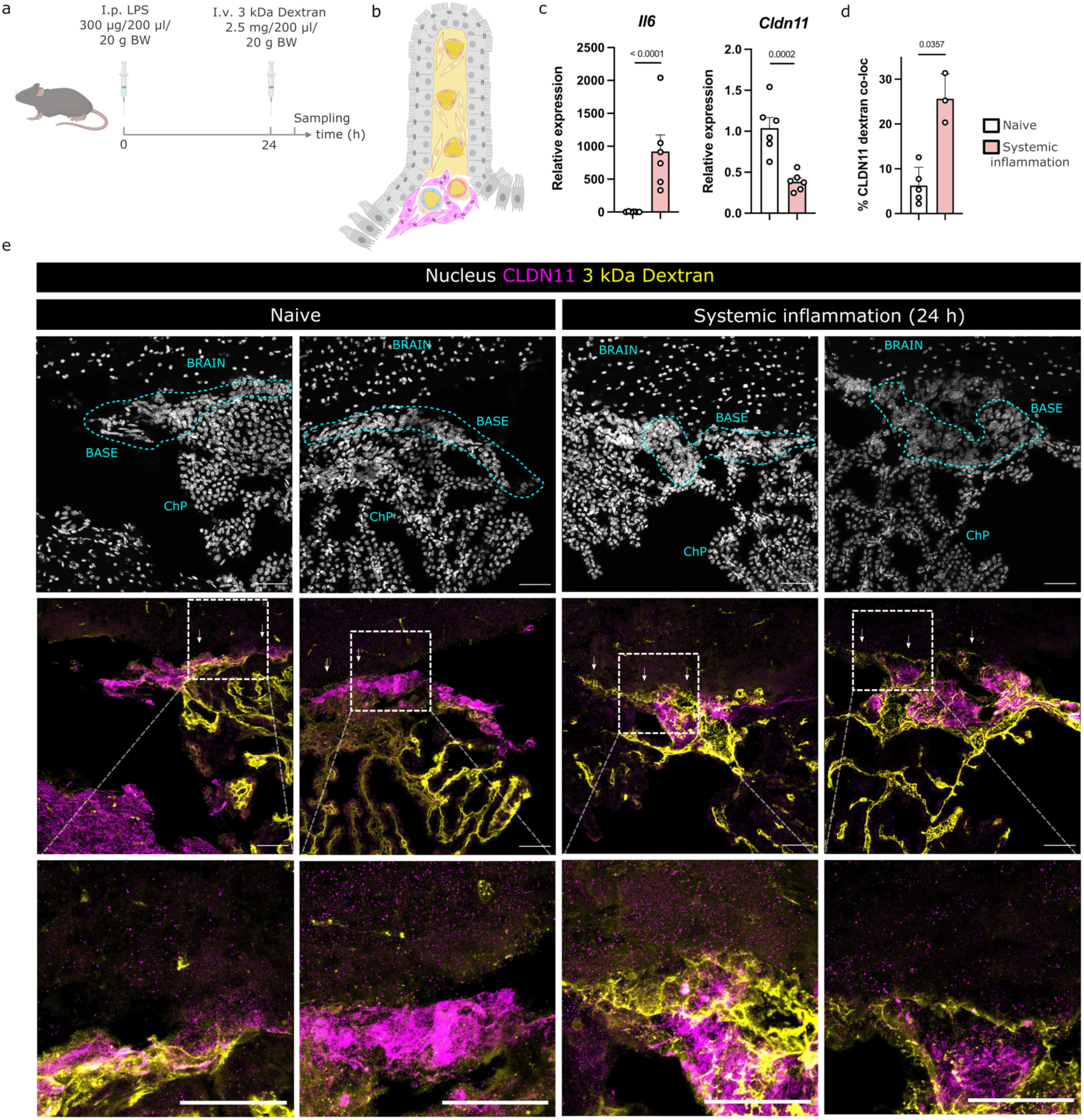
The ChP BBCs are affected by systemic inflammation. (a) Experimental setup of systemic inflammation through i.p. LPS injection. In vivo leakage visualization through tracing of i.v. injected fixable dextrans. (b) Schematic clarifying how i.v. injected dextrans can enter ChP stroma through fenestrated capillaries, but are prevented by ChP BBCs to leak towards brain parenchyma. (c) qPCR analyses of lateral and fourth ventricle ChP tissues of 7-week-old naive vs systemically inflamed mice. N=6 mice per group. (d) Quantification of overlap between CLDN11 and dextran signal in fourth ventricle ChP-containing brain cryosections of 7-week-old naive versus systemically inflamed mice. N=5 (naive) vs 3 (systemic inflammation). (e) IF staining for CLDN11 (magenta) and streptavidin (yellow) in fourth ventricle ChP-containing brain cryosections (with ChP base) of 7-week-old naive (left) versus systemically inflamed (right) mice i.v. injected with 3 kDa biotinylated dextran. Top row shows nuclear stain and anatomical annotation. Middle and bottom row shows IF staining. White arrows indicate neighboring brain parenchyma. Scale bar indicates 50 µm. Representative images for N=5 (naive) versus 3 (systemic inflammation). ChP (choroid plexus); LPS (lipopolysaccharide).

### ChP base barrier cells are conserved in humans

The presence of multiple barriers is essential to maintain brain homeostasis and, as such, these cellular barriers are highly conserved. The ABCs, for example, were shown to be evolutionary conserved across many species^40,49–52^. Based on the transcriptional and functional similarities between ABCs and ChP BBCs, we hypothesized the ChP base barrier to be conserved across species as well. A previous report mentioned CLDN11 expression at the base of the developing rat ChP^3^. To expand on the cross species conservation, we sought out to detect the presence of these cells in human ChP tissue. First, we explored the recently published snRNA-Seq dataset of human post mortem ChP (Fig. 7a)^53^. In addition to macrophages, epithelial, glial, neural, ependymal and endothelial cells, Yang et al also identified a heterogenous mix of mesenchymal cells (Fig. 7a and 7b). In this mesenchymal cluster, a split can be observed between possible stromal-like cells expressing ALPL and CDH11 and base barrier-like cells expressing IGFBP6 and CLDN11 (Fig. 7b). In order to investigate the transcriptomic similarities in more detail, we integrated the human snRNA-Seq and mouse scRNA-Seq data of fibroblasts (Fig. 7c). After BBKNN batch-correction, a partial overlap between datasets was observed (Fig. 7d-7g). Strikingly, the sole cluster containing both human and mouse cells shows a ChP BBC-like signature (Fig. 7f and 7g). We wondered if the partial overlap in transcriptome is also reflected in the expression of our previously identified markers and performed a cross-species conserved marker analysis on the overlapping cluster (Suppl. Table 3). The top conserved markers are S100b/S100B, Cpe/CPE, Vim/VIM, Cystm1/CYSTM1 and excitingly also Igfbp6/IGFBP6 (Fig. 7h). Moreover, we were able to confirm the in vivo expression of IGFBP6 and CLDN11 in human post mortem brain samples. Similar as in mice, we observed CLDN11+ cells clustering at the base of the ChP. These cells also express high levels of IGFBP6 (Fig. 7i). Altogether, we confirm the presence and transcriptional similarities of ChP base barrier cells in both mice and humans.

**Figure 7.**
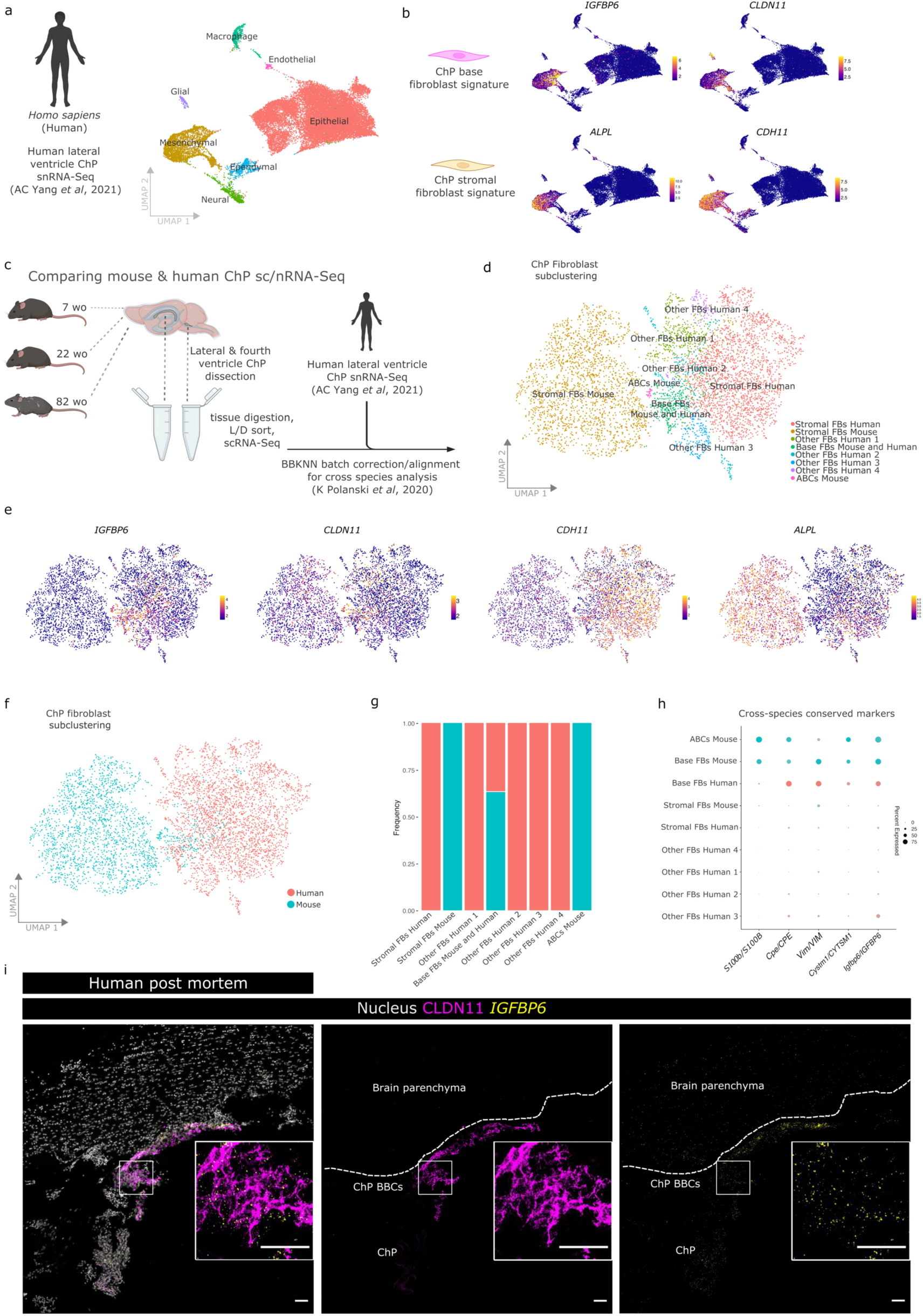
ChP BBCs are conserved in humans and express the same markers. (a) UMAP feature plots showing various cell types detected in snRNA-Seq of human post mortem lateral ventricle ChP tissues described in AC Yang et al, 2021. Pooled data from N=16 patients. (b) UMAP feature plots showing expression of IGFBP6, CLDN11, ALPL and CDH11 by mesenchymal cells in the human ChP. (c) Experimental setup of ChP sampling of 7-, 22- and 82-week-old mice (N=8 pooled mice per age group), scRNA-Seq processing and combination with previously published snRNA-Seq of human post mortem ChP (AC Yang et al, 2021). (d) UMAP feature plots showing sub-clustering of mouse and human ChP fibroblasts only (cell types annotated). (e) UMAP feature plots showing the expression of IGFBP6, CLDN11, CDH11 and ALPL. (f) UMAP feature plots showing sub-clustering of mouse and human ChP fibroblasts only (species annotated). (g) Frequency stacked bar plot showing species contribution to various fibroblast clusters. (h) Dot plot representation of top 5 markers discovered through cross-species conserved marker analysis. (i) RNA FISH and protein multiplex showing expression of IGFBP6 mRNA and CLDN11 protein in human post mortem brain sections with ChP. Zoom in maximum intensity projections of CLDN11 and IGFBP6 shown in squares. Samples shown from a 5-week-old female. Scale bars indicate 50 µm. ChP (choroid plexus); Wo (weeks old); L/D (live/dead); BBKNN (batch balanced k nearest neighbors); FBs (fibroblasts).

## CONCLUSION

Here, we describe and characterize a previously unrecognized barrier located at the base of the ChP. We show that fibroblasts in the ChP represent a heterogeneous population residing in either the stroma or the base of the ChP, and we describe markers to discriminate them from other CNS cells. This ChP base region is in close contact with the SAS and meninges and its unique fibroblast population expresses tight junctions and clusters together thereby sealing the ChP stroma. This way, it provides functional compartmentalization of the ChP, brain and CSF. The ChP base barrier cells (ChP BBCs) are derived from early meningeal mesenchymal precursors, are maintained throughout life and are conserved across species. We show that inflammatory insults affect the ChP base barrier and this results in leakage from blood into the brain. Further investigations into the ChP base barrier in disease will result in novel insights into brain and CNS pathology.

## SUPPLEMENTARY DATA

Suppl Movie 1: movie of ChP base barrier cells (BBCs) located at the ChP-meninges continuity via CLEM, related to figure 4. Adult naive mice were injected with ConA-FITC (cyan) to label blood vessel lumen. Brains were sampled and vibratome sections were stained for CLDN11 (magenta). Confocal microscopy images and SBF-SEM images of same region were overlapped. Note CLDN11+ cell clustering and unique nucleus appearance. **Movies will be made available after peer review and publication.**

Suppl Movie 2: movie of 3D rendering of ChP base (red), arterioles (yellow), venules (cyan) and ChP base barrier cells (BBCs) nuclei (magenta) via SBF-SEM, related to figure 5. Note plug-like shape of clustering ChP BBCs that seal stroma of the ChP from brain parenchyma and CSF. **Movies will be made available after peer review and publication.**

Suppl. Fig. 1 The mesenchymal stem cell marker PDGFRA is expressed by various cells in and nearby the ChP.

Suppl. Fig. 2 ChP base fibroblasts express low levels of ECM genes and produce little collagen.

Suppl. Fig 3 ChP base fibroblasts have a distinct nucleus.

Suppl. Table 1:

1. RNAmarkersList_Original_paper_annotation_Merge_FB_origin_subset shows the differential marker analysis on CNS fibroblast subtypes. See extended materials and methods for details.

Suppl. Table 2:

2. RNAmarkers_and_EnrichGO_ChP_FBs_Merge_FB_origin_subset shows Gene Ontology Enrichment Analysis (GOEA) performed on differentially expressed genes between ChP stromal and base fibroblasts (Suppl. Table 2 and Fig. 4a). Biological Pathway (BP) and Cellular Compartment (CC). See extended materials and methods for details.

Suppl. Table 3:

3. Conserved_ChP_BBC_markers_BBKNN_integrated_Human_and_Mouse shows a cross-species conserved marker analysis on the overlapping cluster containing mouse and human ChP base-barrier like cells. See extended materials and methods for details.

## MATERIALS & METHODS

### Mice and reagents

C57BL/6J mice (Mus musculus) and B6.129S4-Pdgfra^tm11(EGFP)Sor^/J ^54^ were maintained in the VIB Center for Inflammation Research animal facility. Mice were housed in individually ventilated cages in specific pathogen-free conditions, under a 12 h dark/light cycle with food and water ad libitum. We used a mix of male and female mice of 7 to 12 weeks old, unless specified elsewhere. Animal studies were approved by the ethical committee of Ghent University Center for Inflammation Research, Belgium. For the induction of systemic inflammation, we used male and female C57BL/6 J mice which received i.p. injection of 300 µg/200 µl/20 g BW lipopolysaccharide. For in vivo blood vessel tracing, we used male and female 7- to 12-week-old C57BL/6J mice which received i.v. injection of 600 µg/200 µl/20 g BW ConcanavalinA-FITC (Vector labs, VEC.FL-1001) diluted in phosphate-buffered saline solution. For in vivo tracer leakage, we used naive and systemically inflamed male and female 7- to 12-week-old C57BL/6J mice which received either i.v. or i.c.v. injection of Dextran-3000MW-Biotin-Lysin (ThermoFisher, D7135). I.v. dose was 2.5 mg/200 µl/20 g body weight. For i.c.v. injections, mice were first anesthetized with isoflurane and mounted on a stereotactic frame. The body temperature of the mice was maintained at 37 °C using a heating pad. A volume of 5 μl of 100 mg/ml was injected at a rate of 1 µl/min into the left lateral cerebral ventricle with a Hamilton needle, for which the injection coordinates were measured from the bregma (anteroposterior 0.07 cm, mediolateral 0.1 cm, dorsoventral 0.2 cm) using the Franklin and Paxinos mouse brain atlas. Embryonic C57BL/6J mouse work was performed in accordance with the IACUC and relevant guidelines of Boston Children’s Hospital and Masaryk University. Embryonic Tbx18^tm3.1(cre/ERT2)Sev/J 30^x Ai14fl/fl (Gt(ROSA)26Sor^tm14(CAG-tdTomato)Hze^ mice and Mesp1^tm(cre)Ysa^ ^28^x Ai14fl/fl (Gt(ROSA)26Sor^tm14(CAG-tdTomato)Hze^ experiments were performed in accordance with AALAC and were handled in accordance with protocols approved by the University of Colorado Anschutz Medical Campus IACUC committee. Breeding, mice setups and tamoxifen administration were performed as described earlier ^31^.

### Human samples

Human samples were obtained under an IRB approved protocol at Boston Children’s Hospital. A neuropathologist (H.G.W.L.) reviewed the human tissue specimens and recorded non-identifying information. Pictures shown in manuscript are from a 5-week-old female.

### Tissue processing and immunostainings

Mice were sedated with a mix of ketamine (20 mg/ml) and xylazine (4 mg/ml), followed by transcardial perfusion with phosphate-buffered saline (PBS) with 1% Heparin. Brains were dissected and fixed in 4% PFA at 4 °C for 1-4 h, followed by 10-20-30% sucrose gradient, embedding in Neg50-OCT cryogel and storage at -70 °C. Cryosections of 12-18 µm thick were cut using the Cryostar NX70. Sections were then post-fixed with 4% PFA for 10 min, washed with PBS and blocked for 1 h at room temperature with PBS containing 0.1% TritonX100, 0.5% BSA and 2% serum. Serum was matched to species of secondary antibodies. Same blocking buffer was used to dilute primary and secondary antibodies. Primary antibodies were incubated over night at 4 °C, followed by washes in PBS and fluo-conjugated secondary antibody incubation for 2 h at room temperature. Secondary antibodies were used at 1/500 diluted in blocking buffer (Thermofisher: 35503, 35553, A11036, A21094, 35563, 21844). Primary antibodies used in this study: Rabbit anti CLDN11 (1/800, ThermoFisher Scientific, 36-4500); Rat anti CD31 (1/100, BD Pharmigen, 553370); Mouse anti ACTA2-Cy3 (1/400, Sigma Aldrich, C6198); Mouse anti CDH1 (1/200, Becton Dickinson, 610181). After additional washing in PBS and incubation with DAPI for 30 min at room temperature, sections were mounted using PVA-DABCO mounting medium. For the visualization of iv injected Dextran-3000MW-Biotin-Lysin (ThermoFisher, D7135), sections were incubated with Streptavidin-DyLight633 (1/1000, Thermofisher, 21844) diluted in PBS for 30 min at room temperature. Slides were imaged with a Zeiss LSM780 confocal microscope (Carl Zeiss) and analyzed using ImageJ. For co-loc analyses, we used QuPath to calculate the intersect and finally the %CLDN11 signal overlapping with either 3 kDa dextran signal or lineage tracing AI14 reporter signal. Data shown are % co-localization averaged per mouse, with a minimum of 2 sections imaged per ventricle per mouse.

### RNA ISH experiments

For the analyses of Cldn11 and Igfbp6 expression in the developing mouse brain, we investigated the Developing Mouse Brain dataset from the Allen Brain atlas (developingmouse.brain-map.org/). For the analyses of Igfbp6 and Dpep1 expression in adult mouse brains, we performed multiplex RNA-FISH experiments using the Molecular Instruments technology. Tissues were collected as described above, with the sole exception of 24 h fixation in 4% PFA at 4 °C. In brief, cryosections were airdried at room temperature for 15 min, washed in PBS and then baked at 55 °C for 30 min. After additional washing, we briefly incubated Proteinase K 5 µg/ml for 3 min at room temperature, followed by washing and 15 min post-fix in 4% PFA. From this point on, we performed steps as recommended by the company and online available protocol “HCR RNA-FISH protocol for fresh frozen or fixed frozen tissue sections”. We used Dpep1-B4 (NM_007876.2) and Igfbp6-B2 (NM_008344.3) probes. For the analyses of IGFBP6 and CLDN11 in human post mortem FFPE samples, we performed RNAScope analyses. Formalin-Fixed Paraffin-Embedded (FFPE) tissue sections were stained with hematoxylin and eosin (H&E) according to standard methods to identify sections with choroid plexus. Consecutive sections from positively identified sections were used for RNAscope and immunolabeling. Briefly, IGFBP6 mRNA was localized using the RNAscope probe (ACD Bio, Cat No. 496061) and the RNAscope Multiple Fluorescent V2 Assay (Cat. No. 496069). Immediately following the completion of the assay, CLDN11 immunostaining was performed using Rabbit anti-Cldn11 antibody (Thermo, Cat No. 36-4500). Briefly, tissue sections were blocked for 1 hour with 1X PBS + 5% goat serum, followed by incubation with primary antibody at 4 °C overnight and then with Alexa Fluor secondary antibodies (goat anti-Rabbit 594, Thermo Cat. No. A20815), nuclei were counterstained with Hoechst. Slides were imaged with a Zeiss LSM780 confocal microscope (Carl Zeiss) and analyzed using ImageJ.

### Herovici staining

For the analyses of collagens in the ChP, a Herovici staining was performed. In brief, mice were perfused with PBS Heparin, brains isolated and fixed in 4% PFA overnight at 4 °C. Samples were dehydrated through ethanol gradient, xylene and embedded in paraffin. 5 µm thick sections were cut using the HM340E microtome and dried overnight. Samples were rehydrated over xylene and ethanol steps and stained for 10 min with the acid-resistant nuclear staining Weigert’s iron hematoxylin solution (Sigma). After rinsing with tap water for about 5min, samples were incubated with the staining solution (41% picric acid, 4.5% acid fuchsin, 45% Methyl blue, 9% glycerol and 0.45% Lithium carbonate) for 2 min and washed with 1% acetic acid for 2 min. Finally, samples were again dehydrated with ethanol, cleared with xylene and mounted using Entellan^TM^ (Merck Millipore). All steps were performed at room temperature. Slides were imaged with Zeiss AxioScan microscope.

### RNA extraction and RT-qPCR analysis

ChP tissues were dissected as described before ^55^. ChP from perfused mice were snap-frozen in liquid nitrogen and stored at -80 °C. TRIzol (Invitrogen) was used to homogenize the tissue with a the TissueLyser (QIAGEN) in a tube with zirconium oxide beads. Afterwards, chloroform was added and the homogenate was separated in phases by centrifugation at 20.000 g for 15 min at 4 °C. The upper phase was collected and RNA extracted using the Aurum total RNA kit (Bio-Rad) according to the manufacturer’s guidelines. RNA concentration was measured using the Nanodrop-1000 (Thermo Scientific) and reverse-transcribed with the SensiFAST^TM^ cDNA Synthesis Kit (Bioline) to generate cDNA. Primers used: Cldn11 Fw ATGGTAGCCACTTGCCTTCAG; Cldn11 Rev AGTTCGTCCATTTTTCGGCAG; IL6 Fw TAGTCCTTCCTACCCCAATTTCC ; IL6 Rev TTGGTCCTTAGCCACTCCTTC; Rpl Fw CCTGCTGCTCTCAAGGTT; Rpl Rev TGGTTGTCACTGCCTCGTACTT; Ubc Fw AGGTCAAACAGGAAGACAGACGTA; Ubc Rev TCACACCCAAGAACAAGCACA; Gapdh Fw TGAAGCAGGCATCTGAGGG; Gapdh Rev CGAAGGTGGAAGAGTGGGAG; Hprt Fw AGTGTTGGATACAGGCCAGAC; Hprt Rev CGTGATTCAAATCCCTGAAGT. qPCR reaction was performed using the SensiFAST^TM^ SYBR No-ROX Kit (Bioline) on the Roche LightCycler 480 (Applied Biosystems). Data was analyzed using the qbase+ software. Values shown in figures are relative expression values normalized to geometric means of the reference genes.

### Tissue processing and scRNA-Seq

Mice for all groups were first sedated as described above, transcardially perfused with PBS with 1% Heparin after which brain tissue was excised and lateral and fourth ventricle choroid plexus tissue were carefully microdissected and collected in serum-free RPMI at 4 °C. As choroid plexus tissue is small and does not allow for single mouse scRNA-Seq analyses, we pooled choroid plexus tissues from 8 mice to generate single pooled samples for naive 7-week-old (female), 22-week-old (male) and 82-week-old (male) mouse groups, for the lateral and fourth ventricle ChP tissue separately. After tissue collection, samples were pre-cut with small scissors and diluted and digested in 2:3 stock solution of CollagenaseI (10 U/ml, Worthington), Collagenase IV (400 U/ml, Worthington) and DNaseI (30 U/ml, Worthington). Incubation was at 37 °C for 30 min with up/down pipetting every 10 min. Samples were strained through a 70 µm filter (Falcon) and washed with a total of 700 µl RPMI. Samples were centrifuged for 7 min 300 g at 4 °C, supernatant discarded and pellet resuspended in 250 µl FACS buffer (1x HBSS without Ca^2+^/Mg^2+^, 2 mM EDTA, 2% BSA). Single live cells were sorted using the BD FACS ARIAII and loaded onto a Chromium GemCode Single Cell Instrument (10x Genomics) to generate single-cell gel beads in emulsion (GEMs). Libraries were prepared with the GemCode Single Cell 3’ Gel Bead and Library Kit v1 (10x Genomics) and the Chromium i7 Multiplex Kit (10x Genomics) according to manufacturer’s instructions. Afterwards,10X barcoded cDNA was prepared for Illumina HiSeq4000 Next Gen Sequencing. Sequencing was performed at the VIB Nucleomics Core (VIB, Leuven). Demultiplexing of the raw data was done with 10x Cell ranger software (v2.0.1). Obtained reads from demultiplexing were used as input for ‘cellranger count’ to align reads to the human reference genome using STAR and collapse them to unique molecular identifier (UMI) counts. Resulting expression matrix was used for follow-up analyses, which are described in the extended materials and methods section. After processing and doublet removal, we ended with 11.075 cells of pooled LV and 4V ChP samples of 7-week-old mice, 18.783 cells of pooled LV and 4V ChP samples of 22-week-old mice; 18.222 cells of pooled LV and 4V ChP samples of 82-week-old mice. The tissue processing and sc/nRNA-Seq analyses of embryonic and human choroid plexus samples are described in ^11^ and ^56^, respectively.

### Sc/nRNA-Seq merged datasets and analyses

We processed and combined various sc and snRNA-Seq datasets. The details and rationale for the processing pipelines are described in the “Extended materials and methods” section.

### Electron microscopy

Mice for EM analyses were sedated as described above, brains were isolated and fixed in 2% PFA with 2% glutaraldehyde for 1h at room temperature, followed by overnight incubation at 4 °C. After thorough washing with PBS, brains were embedded in 5% agarose and vibratome section of 100-200 µm thick were cut with the Leica Vibratome VT1200S. Sections with clear choroid plexus base region were selected, and further cut into small square tissues. Tissues were processed for SBF-SEM analyses with the “OTO” protocol as described previously ^57^ with the sole exception that we did not use the Spurr’s resin but Embed 812. Samples were mounted onto aluminum pins and trimmed and SBF-SEM imaged with the Zeiss Merlin with 3View2 (Gatan). When desired, regions with clear choroid plexus base region were also transferred to and imaged with the Zeiss Crossbeam 450 for FIB-SEM imaging ^39^. For tracing we used Microscopy Image Browser (MIB) software and 3D rendering was done in Imaris.

### Correlative light electron microscopy

CLEM workflow was performed as described earlier ^38,58,59^. In brief, mice were i.v. injected with 1 mg/200 µl/20 g BW ConcanavalinA-FITC and immediately sedated with a mix of ketamine (20 mg/ml) and xylazine (4 mg/ml). After 5 min, mice were decapitated, brains sampled and fixed in 4% PFA with 0.25% glutaraldehyde for 4 h at 4 °C. Brains were washed with PBS and embedded in 5% agarose and cut into 50 µm-thick vibratome sections using the Leica Vibratome VT1200S. We performed free-floating anti-CLDN11 immunostainings on these sections. After mounting, samples were first scanned for regions of interest (ROIs) and branded using Near Infra-Red Branding using the Zeiss LSM780 with MaiTai laser ^60^. ROIs were then imaged through LSM880 FastAiry super resolution imaging. These same samples were then processed for volume EM as described above. Overlaying both confocal LM and volume EM datasets was done with the Bigwarp plugin of ICY ^61^ and using branching points of the blood vessels as points for landmark-based registration.

### Statistical analyses

Sample sizes for different experiments were based on standard power calculations and previous experience with similar experiments. For most experiments we used a mix of both male and female mice, unless stated otherwise. Statistical analysis was done using Prism9. Dotplots show average value per mouse ± SD. Normal distribution of data was checked using the Kolmogorov-Smirnov test and Shapiro-Wilk tests. For CLDN11-AI14 co-loc analysis, data was normally distributed according to Shapiro-Wilk test and Welch’s t test was performed. For CLDN11-Dextran co-loc analysis, data was not normally distributed and we therefore performed the most strict Mann Whitney test. For qPCR analysis, data was normally distributed after log-transformation and we therefore performed unpaired t-tests. Statistical tests performed on scRNA-Seq data (e.g. through GSEA) were corrected for multiple testing. Details for scRNA-Seq related tests and statistics are described in the extended materials and methods section.

### Data & code availability

All raw sequencing data enclosed in this publication has been deposited in NCBI’s Gene Expression omnibus ^62^ and will be made accessible after peer-review and publication.

## Acknowledgments

We would like to thank the VIB Bioimaging Core & VIB Flow Core for training, support and access to the instrument park. We would like to thank the VIB Single Cell Core for outstanding technical support regarding single-cell omics workflows. We would like to thank the VIB Tech Watch for early access grants. This work was supported by the Research Foundation-Flanders (FWO PhD fellowships (11D0520N, 11A6420N, 1157621N), Postdoctoral junior fellowship (1268823N) and FWO Junior research project (G055121N)) and the Baillet Latour Grant for Medical Research. Figures were made with BioRender.com.

---

**Supplementary Figure 1.**
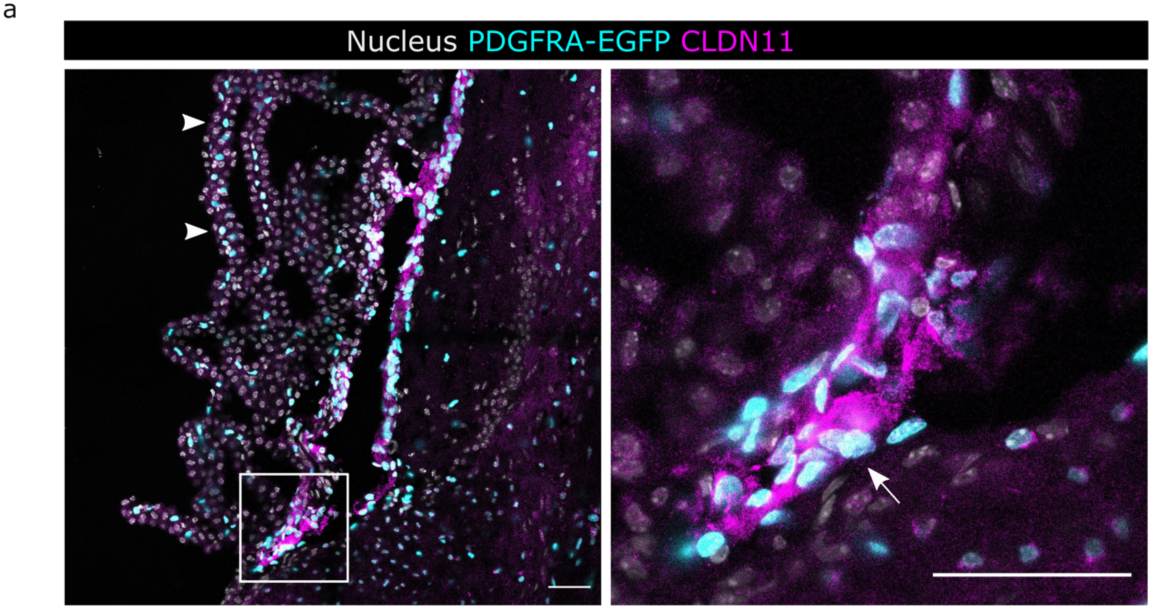
The mesenchymal stem cell marker PDGFRα is expressed by various cells in and nearby the ChP. (a) IF staining for CLDN11 (magenta) in third ventricle ChP-containing brain cryosections of 7-week-old Pdgfra-H2B-eGFP transgenic mice. White arrow: ChP base fibroblasts. White arrowhead: ChP stromal cells. Scale bar indicates 50 µm. Representative image for N=2 mice.

**Supplementary Figure 2.**
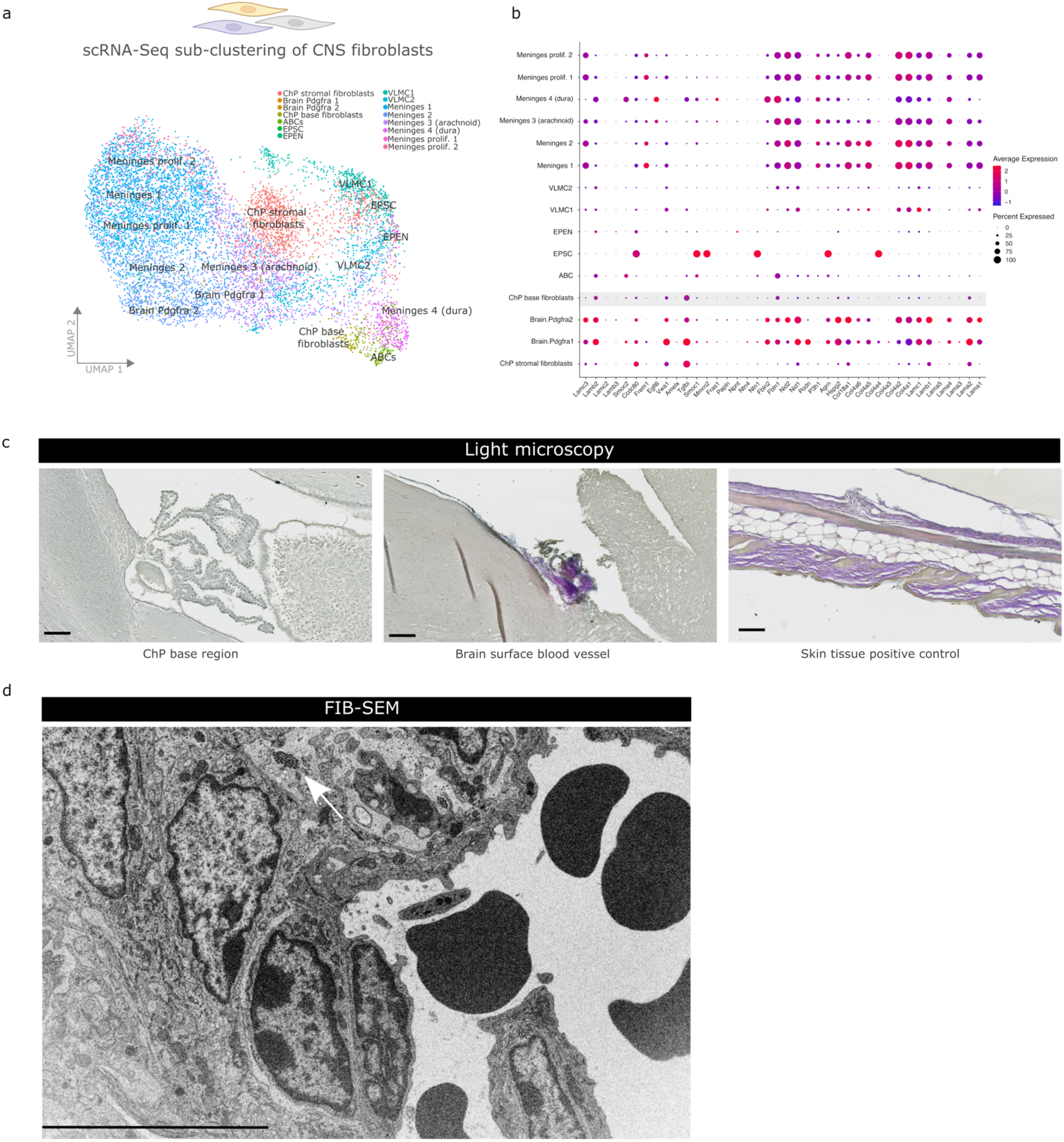
ChP base fibroblasts express low levels of ECM genes and produce little collagen. (a) UMAP visualization showing scRNA-Seq data of CNS fibroblasts subclustering. (b) Dot plot representation showing expression of selected ECM-genes by CNS fibroblasts. (c) Herovici staining to discriminate mature (red) and immature (blue) collagens in brain sections with ChP and control samples. Scale bar indicates 50 µm. Representative image for N=2 mice. (d) FIB-SEM analysis on brain vibratome sections with ChP base region. White arrow indicates only small collagen bundles nearby ChP base fibroblasts. Scale bar indicates 10 µm. Data from single mouse. ChP (choroid plexus); VECC (vascular endothelial cells capillary); VSMCA (vascular smooth muscle cells arterial); VAC (Vascular associated cells/pericytes); NSC (neuronal stem cells); VLMC (vascular leptomeningeal cells); EPSC (ependymal cells spinal cord); ABC (arachnoid barrier cells); FIB-SEM (focused-ion-beam scanning electron microscopy).

**Supplementary Figure 3.**
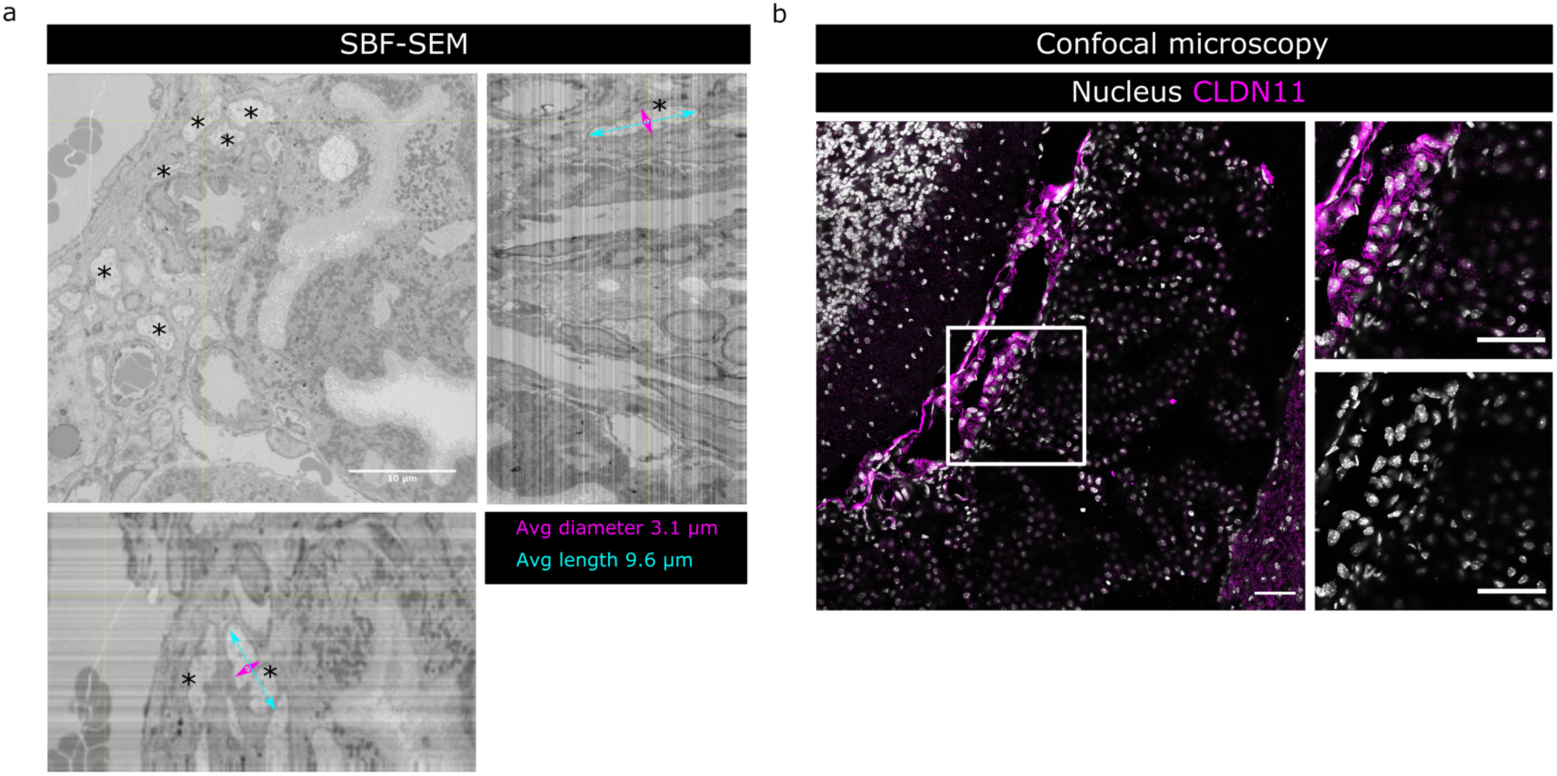
ChP base fibroblasts have a distinct nucleus. (a) SBF-SEM analyses on fourth ventricle ChP-containing vibratome sections (different planes shown) of 7-week-old mice. Indicated are the average diameter and length of the ChP base fibroblasts (N=13 cells). Note elongated shape, pale nucleus with dense heterochromatin edge. Scale bar indicates 10 µm. Data from single mouse. (b) IF staining for CLDN11 (magenta) in fourth ventricle ChP-containing brain sections of 7-week-old mice. Note elongated nuclei of CLDN11+ cells that cluster together. Scale bar indicates 50 µm unless specified differently. Data representative for N=4 mice.

## REFERENCES

1. Marchetti, L. & Engelhardt, B. Immune cell trafficking across the blood-brain barrier in the absence and presence of neuroinflammation. Vasc. Biol. (Bristol, England) 2, H1–H18 (2020).

2. Chow, B. W. & Gu, C. The molecular constituents of the blood-brain barrier. Trends Neurosci. 38, 598 (2015).

3. Weller, R. O., Sharp, M. M., Christodoulides, M., Carare, R. O. & Møllgård, K. The meninges as barriers and facilitators for the movement of fluid, cells and pathogens related to the rodent and human CNS. Acta Neuropathologica 135, 363–385 (2018).

4. Ghersi-Egea, J.-F. et al. Molecular anatomy and functions of the choroidal blood-cerebrospinal fluid barrier in health and disease. Acta Neuropathol. 135, 337–361 (2018).

5. Wilting, J. & Christ, B. An experimental and ultrastructural study on the development of the avian choroid plexus. Cell Tissue Res. 1989 2553 255, 487–494 (1989).

6. Dorrier, C. E., Jones, H. E., Pintarić, L., Siegenthaler, J. A. & Daneman, R. Emerging roles for CNS fibroblasts in health, injury and disease. Nature Reviews Neuroscience 1–12 (2021). doi:10.1038/s41583-021-00525-w

7. Derk, J., Jones, H. E., Como, C., Pawlikowski, B. & Siegenthaler, J. A. Living on the Edge of the CNS: Meninges Cell Diversity in Health and Disease. Front. Cell. Neurosci. 0, 245 (2021).

8. McLone, D. G. & Bondareff, W. Developmental morphology of the subarachnoid space and contiguous structures in the mouse. Am. J. Anat. 142, 273–293 (1975).

9. Dasgupta, K. & Jeong, J. Developmental Biology of the Meninges. Genesis 57, e23288 (2019).

10. DeSisto, J. et al. Single-Cell Transcriptomic Analyses of the Developing Meninges Reveal Meningeal Fibroblast Diversity and Function. Dev. Cell 54, 43–59.e4 (2020).

11. Dani, N. et al. A cellular and spatial map of the choroid plexus across brain ventricles and ages. Cell 184, 3056–3074.e21 (2021).

12. Shuangshoti, S. & Netsky, M. G. Histogenesis of choroid plexus in man. Am. J. Anat. 118, 283–315 (1966).

13. Greiner, T. et al. Morphology of the murine choroid plexus: Attachment regions and spatial relation to the subarachnoid space. Front. Neuroanat. 16, (2022).

14. Van Hove, H. et al. A single-cell atlas of mouse brain macrophages reveals unique transcriptional identities shaped by ontogeny and tissue environment. Nat. Neurosci. 22, 1021–1035 (2019).

15. Vanlandewijck, M. et al. A molecular atlas of cell types and zonation in the brain vasculature. Nature 554, 475–480 (2018).

16. PA Johansson, M Irmler, D Acampora, J Beckers, A Simeone, M. G. The transcription factor Otx2 regulates choroid plexus development and function. Development 140, 1055–1066 (2013).

17. Zeisel, A. et al. Molecular Architecture of the Mouse Nervous System Resource Molecular Architecture of the Mouse Nervous System. Cell 174, 999–1014 (2018).

18. Shah, P. T. et al. Single-Cell Transcriptomics and Fate Mapping of Ependymal Cells Reveals an Absence of Neural Stem Cell Function. Cell 173, 1045–1057.e9 (2018).

19. Cougnoux, A. et al. Single Cell Transcriptome Analysis of Niemann–Pick Disease, Type C1 Cerebella. Int. J. Mol. Sci. 21, 1–23 (2020).

20. Vidovic, D., Davila, R. A., Gronostajski, R. M., Harvey, T. J. & Piper, M. Transcriptional regulation of ependymal cell maturation within the postnatal brain. Neural Dev. 13, 1–9 (2018).

21. Saurav Roy Choudhury, A., et al. Dipeptidase-1 Is an Adhesion Receptor for Neutrophil Recruitment in Lungs and Liver. Cell 178, 1205–1221.e17 (2019).

22. Lau, A. et al. Dipeptidase-1 governs renal inflammation during ischemia reperfusion injury. Sci. Adv. 8, (2022).

23. Bach, L. A. Recent insights into the actions of IGFBP-6. J. Cell Commun. Signal. 9, 189–200 (2015).

24. Gama Sosa, M. A., De Gasperi, R., Perez, G. M., Hof, P. R. & Elder, G. A. Hemovasculogenic origin of blood vessels in the developing mouse brain. J. Comp. Neurol. 529, 340–366 (2021).

25. Hu, H., Lin, S., Wang, S. & Chen, X. The Role of Transcription Factor 21 in Epicardial Cell Differentiation and the Development of Coronary Heart Disease. Front. Cell Dev. Biol. 8, 457 (2020).

26. Plotkin, M. & Mudunuri, V. Pod1 induces myofibroblast differentiation in mesenchymal progenitor cells from mouse kidney. J. Cell. Biochem. 103, 675– 690 (2008).

27. Farahani, R. M. & Xaymardan, M. Platelet-Derived Growth Factor Receptor Alpha as a Marker of Mesenchymal Stem Cells in Development and Stem Cell Biology. Stem Cells Int. 2015, (2015).

28. Saga, Y. et al. MesP1 is expressed in the heart precursor cells and required for the formation of a single heart tube. Development 126, 3437–3447 (1999).

29. Yoshida, T., Vivatbutsiri, P., Morriss-Kay, G., Saga, Y. & Iseki, S. Cell lineage in mammalian craniofacial mesenchyme. Mech. Dev. 125, 797–808 (2008).

30. Guimarães-Camboa, N. et al. Pericytes of Multiple Organs Do Not Behave as Mesenchymal Stem Cells In Vivo. Cell Stem Cell 20, 345–359.e5 (2017).

31. Derk, J. et al. Formation and Function of the Meninges Arachnoid Barrier Around the Developing Brain. SSRN Electron. J. (2022). doi:10.2139/SSRN.4143787

32. Kelly, K. K. et al. Col1a1+ perivascular cells in the brain are a source of retinoic acid following stroke. BMC Neurosci. 17, (2016).

33. Bonnans, C., Chou, J. & Werb, Z. Remodelling the extracellular matrix in development and disease. Nat. Rev. Mol. Cell Biol. 2014 1512 15, 786–801 (2014).

34. Kanchanawong, P. & Calderwood, D. A. Organization, dynamics and mechanoregulation of integrin-mediated cell–ECM adhesions. Nat. Rev. Mol. Cell Biol. 2022 1–20 (2022). doi:10.1038/s41580-022-00531-5

35. Talbot, J. C. et al. Pharyngeal morphogenesis requires fras1-itga8-dependent epithelial-mesenchymal interaction. Dev. Biol. 416, 136–148 (2016).

36. Marek, I., Hilgers, K. F., Rascher, W., Woelfle, J. & Hartner, A. A role for the alpha-8 integrin chain (itga8) in glomerular homeostasis of the kidney. Mol. Cell. Pediatr. 2020 71 7, 1–4 (2020).

37. Ji, Q. et al. Primary tumors release ITGBL1-rich extracellular vesicles to promote distal metastatic tumor growth through fibroblast-niche formation. Nat. Commun. 11, (2020).

38. Kremer, A. et al. A workflow for 3D-CLEM investigating liver tissue. J. Microsc. 281, 231–242 (2021).

39. Vanslembrouck, B. et al. Three-dimensional reconstruction of the intercalated disc including the intercellular junctions by applying volume scanning electron microscopy. Histochem. Cell Biol. 149, 479–490 (2018).

40. Nabeshima, S., Reese, T. S., Landis, D. M. D. & Brightman, M. W. Junctions in the meninges and marginal glia. J. Comp. Neurol. 164, 127–169 (1975).

41. Saboori, P. & Sadegh, A. Histology and Morphology of the Brain Subarachnoid Trabeculae. Anat. Res. Int. 2015, 1–9 (2015).

42. Balin, B. J., Broadwell, R. D., Salcman, M. & El-Kalliny, M. Avenues for entry of peripherally administered protein to the central nervous system in mouse, rat, and squirrel monkey. J. Comp. Neurol. 251, 260–280 (1986).

43. Galea, I. The blood–brain barrier in systemic infection and inflammation. Cell. Mol. Immunol. 2021 1811 18, 2489–2501 (2021).

44. Vandenbroucke, R. E. et al. Matrix metalloprotease 8-dependent extracellular matrix cleavage at the blood-CSF barrier contributes to lethality during systemic inflammatory diseases. J. Neurosci. 32, 9805–16 (2012).

45. Mottahedin, A., Smith, P. L. P., Hagberg, H., Ek, C. J. & Mallard, C. TLR2-mediated leukocyte trafficking to the developing brain. J. Leukoc. Biol. 101, 297–305 (2017).

46. Mottahedin, A. et al. N-acetylcysteine inhibits bacterial lipopeptide-mediated neutrophil transmigration through the choroid plexus in the developing brain. Acta Neuropathol. Commun. 8, 4 (2020).

47. Reboldi, A. et al. C-C chemokine receptor 6–regulated entry of TH-17 cells into the CNS through the choroid plexus is required for the initiation of EAE. Nat. Immunol. 10, 514–523 (2009).

48. Cui, J. et al. Inflammation of the Embryonic Choroid Plexus Barrier following Maternal Immune Activation. Dev. Cell 0, (2020).

49. Nakao, T., Ishizawa, A. & Ogawa, R. Observations of vascularization in the spinal cord of mouse embryos, with special reference to development of boundary membranes and perivascular spaces. Anat. Rec. 221, 663–677 (1988).

50. Vandenabeele, F., Creemers, J. & Lambrichts, I. Ultrastructure of the human spinal arachnoid mater and dura mater. J. Anat. 189, 417 (1996).

51. Rascher, G. & Wolburg, H. The tight junctions of the leptomeningeal blood-cerebrospinal fluid barrier during development. J. Hirnforsch. 38, 525–540 (1997).

52. Brøchner, C. B., Holst, C. B. & Møllgård, K. Outer brain barriers in rat and human development. Front. Neurosci. 0, 75 (2015).

53. Yang, A. C. et al. Dysregulation of brain and choroid plexus cell types in severe COVID-19. Nat. 2021 5957868 595, 565–571 (2021).

54. Hamilton, T. G., Klinghoffer, R. A., Corrin, P. D. & Soriano, P. Evolutionary divergence of platelet-derived growth factor alpha receptor signaling mechanisms. Mol. Cell. Biol. 23, 4013–4025 (2003).

55. Van Wonterghem, E. et al. Microdissection and Whole Mount Scanning Electron Microscopy Visualization of Mouse Choroid Plexus. J. Vis. Exp. 2022, (2022).

56. Yang, A. C. et al. A human brain vascular atlas reveals diverse mediators of Alzheimer’s risk. Nat. 2022 1–8 (2022). doi:10.1038/s41586-021-04369-3

57. Steeland, S. et al. Counteracting the effects of TNF receptor-1 has therapeutic potential in Alzheimer’s disease. EMBO Mol. Med. 10, (2018).

58. Bonnardel, J., et al. Stellate Cells, Hepatocytes, and Endothelial Cells Imprint the Kupffer Cell Identity on Monocytes Colonizing the Liver Macrophage Niche. Immunity 51, 638–654.e9 (2019).

59. Kremer, A., Lippens, S. & Guérin, C. J. Correlative Light and Electron Microscopy: Methods and Applications. eLS 1–10 (2016). doi:10.1002/9780470015902.A0025983

60. Bishop, D. et al. Near-infrared branding efficiently correlates light and electron microscopy. Nat. Methods 8, 568–572 (2011).

61. Bogovic, J. A., Hanslovsky, P., Wong, A. & Saalfeld, S. Robust registration of calcium images by learned contrast synthesis. Proc. - Int. Symp. Biomed. Imaging 2016-June, 1123–1126 (2016).

62. Edgar, R., Domrachev, M. & Lash, A. E. Gene Expression Omnibus: NCBI gene expression and hybridization array data repository. Nucleic Acids Res. 30, 207–210 (2002).

