## Extended Materials and Methods for "Newly discovered base barrier cells provide compartmentalization of choroid plexus, brain and CSF"

### Extended material and methods

**Custom code availability:**
Our custom pipelines will be made available our GitHub after peer review and publication.

**7-week-old ChP LV & 4V aggregate object**
The aggregation of two samples, 7-week-old mice ChP LV and 4V datasets, was done using 'cellranger aggr'. The post-normalization mean reads per cell of the aggregate object was 15,100 reads, as calculated by Cell Ranger. The data were pre-processed using the scater R package (v1.14.6) according to the workflow proposed by Lun and colleagues [1]. Outlier cells were identified based on three metrics (library size, number of expressed genes and mitochondrial proportion) and cells were tagged as outliers according to median absolute deviation (MAD), respectively four MADs for library size and gene count and five MADs for the mitochondrial proportion. Some additional cells were removed which still appeared as outliers based on PCA. Low-abundance genes were removed during the Seurat object creation. Log normalization of the raw counts, highly variable gene detection according to the vst method, data scaling, principal component analysis (PCA), unsupervised clustering and the dimensionality reduction plot creation were done according to the documentation from Seurat (v3.1.4) [2]. Twenty-five PCA dimensions and a resolution of 0.8 were used to create the final Uniform Manifold Approximation and Projection (UMAP) dimensionality reduction plot and clustering. Doublets and low-quality cells were removed based on DoubletFinder (v2.0.2) prediction results and quality metrics [3]. Some additional manual clustering was performed to split up certain clusters based on marker expression. Marker genes per identified subpopulation were found using the FindAllMarkers and FindMarkers functions of the Seurat package. The final object shown in this paper contains 11,383 cells and 8 annotated clusters.

**Fibroblast origin complete object**
To investigate the ontogeny of the Type II Fibroblasts we integrated some of our scRNA-Seq datasets with public scRNA-Seq datasets containing various ependymal, vascular and mesenchymal cell types from the brain.

| Name | Cell type | Age | Species | Origin/Paper |
|---|---|---|---|---|
| GSM_2677817 | Ependymal cells | 8w | Mouse | Shah et al. [4] |
| GSM_2677818 | Ependymal cells | 8w | Mouse | Shah et al. [4] |
| GSM_2677819 | Ependymal cells | 8w | Mouse | Shah et al. [4] |
| GSE98816 | Brain vascular cells | 10-19w | Mouse | Vanlandewijck et al. [5] |
| ChP 4V_7w<br>ChP LV_7w<br>ChP_4V_22w<br>ChP_LV_22w | ChP vascular associated cells | 7w<br>7w<br>22w<br>22w | Mouse | Verhaege et al. |
| GSE150219 | Meningeal fibroblasts | E14 | Mouse | Desisto et al. [6] |
| l6_r3_vascular_cells | CNS vascular cells | 6-8w | Mouse | Zeisel et al. [7] |
| l6_r4_ependymal_cells | Ependymal cells | 6-8w | Mouse | Zeisel et al. [7] |

For our datasets and the public datasets which provided only the raw counts, the datasets were pre-processed using the scater R package (v1.14.6). Outlier cells were again identified based on the MAD of the same three metrics, now respectively three MADs for each metric. No additional PCA cell removal was performed. Low-abundance genes were removed during the Seurat object creation. Log normalization of the raw counts was performed and then each Seurat object was saved. For the GSE150219 dataset from the Desisto paper [6] normalized counts and metadata were provided after filtering. No further pre-processing was needed. A Seurat object was created from the provided data. For the Zeisel datasets [7] we explored their Mouse brain atlas available at http://mousebrain.org/. We downloaded two loom files, vascular cells at taxonomy level 3 and ependymal cells at taxonomy level 4. These datasets were clustered by taxonomy after all filtering and sorting steps. This means no further pre-processing was needed. The loom files just had to be converted to Seurat objects. The data collection and pre-processing resulted in 8 individual Seurat objects. These objects with the normalized data were merged together using Seurat. It was determined previously in other analyses that this workflow achieved similar results as the CellRanger aggregate method. For the merged object, the rest of the previously described workflow (after log normalization) was run with one addition. After the PCA step, Harmony (v1.0) batch correction [8] with a standard theta value of 2 was performed to integrate the various samples with each other. This resulted in a mixing of the different samples across the cell types, so it was deemed that the samples were sufficiently integrated with Harmony. Forty Harmony corrected PCA dimensions and a resolution of 0.8 were used to create the final UMAP plot and clustering. Some minor additional manual clustering was performed to split up one cluster based on marker expression and the metadata of the individual objects was used to inform the annotation process. Subsequently, a subset was made without the immune cell clusters and Choroid Plexus Epithelial Cell clusters because they were not relevant to the analysis. Outlier cells which clustered by these clusters were also removed. The final object contains 29,394 cells and 8 annotated clusters.

**Fibroblast origin subset object**

Following this initial analysis to confirm the fibroblast origin of our Type II Fibroblasts, we created a further subset with only the Fibroblast cluster (cluster idents 1, 9 and 16) to accurately compare the various fibroblasts across the datasets. After the subset, the same workflow was performed as with the full merged object above (including Harmony batch correction). Thirty Harmony corrected PCA dimensions and a resolution of 1 were used to create the final UMAP plot and clustering. With the higher resolution some extra contaminating cells were identified and removed. The final object contains 9,128 cells and 6 newly annotated clusters. When the cleaned up old annotation from the individual objects is used, then there are 15 annotated clusters.

**Marker analysis dot plot Fibroblast origin subset object**

Marker analysis via the Seurat function FindAllMarkers was performed using the cleaned up old annotation which stems from the original annotation of the individual datasets. We wanted to find markers which were highly expressed in the various clusters, so the min.pct parameter was set at a very strict level of 0.6 to avoid lowly expressed markers. Next a filtering step was performed on this marker list to remove contaminating Choroid Plexus

Epithelial (ChPE) cell markers from our two fibroblast populations (utilizing the ChPE marker list acquired from the original merged object with all cell types). This resulted in a strict filtered marker list which was sorted according to the score column. The top 3 markers for each cluster were visualized in a dot plot.

**Functional annotation Fibroblast origin subset object**

Gene ontology enrichment analysis was performed using the clusterProfiler R package (v3.14.3). Two ontologies ("Biological Pathway" and "Cellular Compartment") were included and a p-value cut-off of 0.05 was utilized. All the genes from our scRNA-Seq Seurat object were used as the background genes for the analysis. This was conducted on the marker gene sets for ChP Stromal and Base Fibroblasts. These gene sets were acquired by running the FindMarker function between Stromal and ChP base Fibroblasts in the final Fibroblast origin subset object utilizing the original individual annotation. The resulting gene set was split up based on logFC, positive logFC genes were linked to ChP Stromal Fibroblasts and negative logFC genes to ChP Base Fibroblasts. GO enrichment analysis was run on the two respective gene lists. The results were combined into one dot plot by running the merge_result function from clusterProfiler. The top 25 significantly enriched GO categories per Ontology and per Cell type are featured in the dot plot (due to overlap in top GO categories for BP and the limited number of GO categories for CC this amounts to less than 50 categories each). These top GO categories are ordered according to adjusted p-value, prioritizing Stromal Fibroblasts. The adjusted p-value is displayed as the color of the dot and the size of the dot is determined by the GeneRatio parameter, which is the ratio of the input DE gene set annotated in the respective GO term.

**ChP Fibroblast age object**

To analyze how the transcriptome of fibroblasts varies with age, we integrated various ChP scRNA-Seq samples of differently aged mice: embryonal, 7 weeks old, 22 weeks old and 82 weeks old. The 7-, 22- and 82-week-old mice ChP LV and 4V datasets were our own datasets. These six datasets were analyzed individually according to the pipeline described previously and the annotation was used to subset the fibroblast cells from the raw data of each dataset. Thereafter, all these fibroblast cells were combined into one Seurat object. The ChP scRNA-seq embryonal data from the Lehtinen lab [9] was acquired via the Single Cell Portal from the Broad Institute (https://singlecell.broadinstitute.org/single_cell/study/SCP1365/choroid-plexus-cell-atlas). The raw and normalized counts were provided together with some metadata and dimensionality reduction coordinates. This was used to create a Seurat object which was then subsequently subsetted to only include the mesenchymal cells. Lastly, the 3$^{rd}$ ventricle cells were removed from the object because our datasets do not include the 3$^{rd}$ ventricle Following this initial prep, these two Seurat objects were integrated via Canonical Correlation Analysis (CCA) according to the scRNA-Seq integration procedure from Seurat with the functions SelectIntegrationFeatures, FindIntegrationAnchors and IntegrateData (https://satijalab.org/seurat/articles/integration_introduction.html) using default parameters. This resulted in an integrated data assay which was used as the default assay for the rest of the Seurat workflow as described previously. One exception though was the visualization of the expression of typical cell type markers using violin plots and feature plots where the standard log-normalized RNA assay was used. Thirty integrated PCA dimensions and a resolution of 0.8 were used to create the UMAP plot and clustering. Among our initially selected fibroblast cells there were however some contaminating cells which now clustered separately and were removed. The final object contains 4,257 cells and 7 annotated clusters.

**Stacked bar chart ChP Fibroblast age object**

The contribution from the various ages to the two fibroblast populations of interest was analyzed and visualized with a percentage stacked bar chart. The differences in total cells between the four ages was taken into account by comparing relative amounts for each cell population instead of absolute numbers.

**ChP Fibroblast species object**

To check the cross-species nature of our mouse ChP Base Fibroblasts, we looked for a human dataset of the Choroid Plexus containing a sizeable mesenchymal cell proportion. The snRNA-seq dataset from the Yang et al. [10] study fit the bill. The human Choroid Plexus Seurat object was acquired through the Stanford Shiny app (https://twc-stanford.shinyapps.io/scrna_brain_covid19/) as a rds file. This was the analyzed object with all the metadata, so no pre-processing was needed. Applicable human fibroblast markers were visualized to confirm the identity of the mesenchymal cell cluster to then subset the object to only include the mesenchymal cells to integrate with our mouse datasets. For this analysis we used the same six datasets from our lab which we used in the previous Fibroblast age object analysis, but the merged object was now first analyzed completely on its own so that contaminating cells could be removed before integration. Integrating the merged object from our 6 mouse datasets with the mesenchymal subset of the human dataset proved to be a challenge. As we were integrating scRNA-seq with snRNA-seq across species, it was necessary to apply a stronger batch effect correction method than before. After studying a recent benchmark of integration methods [8], we chose to implement the batch-balanced k nearest neighbors (BBKNN) algorithm [11] to achieve an optimal result. Both Seurat objects were updated through the UpdateSeuratObject function to be able to analyze it further in a newer R environment (v4.0.5) and python environment (v3.9.9) for BBKNN (v.1.5.1). Before integration via BBKNN, the gene symbols of the human mesenchymal subset first had to be converted to their mouse orthologs. This was accomplished by utilizing the Ensembl database through the biomaRt package (v2.46.3). Human genes without mouse orthologs were removed from the count matrix. A new human Seurat object was created from this converted raw count matrix. Metadata from the original object was transferred to this new human Seurat object. When merging the human mesenchymal dataset with our merged mouse dataset, the count matrices were subsetted to the common genes between the two datasets. Subsequently, a part of the previously described workflow was run on the merged cross-species object: log-normalization was applied to the raw counts, highly variable genes were determined, the normalized counts were scaled and PCA was run in R. Thereafter, the cross-species Seurat object was converted to an AnnData python object to run BBKNN in python. Before running BBKNN, ridge regression was used with the species metadata as the technical effect and the initial leiden clustering as the biological grouping, which should improve the integration. BBKNN was then run on the first 30 PCA dimensions. The new integrated PCA dimensions were then used to determine a new clustering (resolution 0.5) and the UMAP dimensionality reduction. After the python analysis was completed, the AnnData object was converted back into a Seurat object for further downstream analysis in R. The final object contains 5,406 cells and 8 annotated clusters.

**Stacked bar chart ChP Fibroblast species object**
The contribution from the two species to the 8 annotated fibroblast clusters was analyzed and visualized with a percentage stacked bar chart. The difference in total cells between the two datasets was taken into account by comparing relative amounts for each cell population instead of absolute numbers.

**Conserved marker analysis dot plot ChP Fibroblast species object**
To determine the markers which are conserved between human and mouse for the ChP Base Fibroblast population, the FindConservedMarkers function from Seurat was run on the ChP Base FBs cluster (versus all other cells) with species as the grouping variable and otherwise default parameters. This returned a ranked list of putative conserved markers with the associated statistics for each species and a couple of summary columns too. This list of conserved markers was ordered according to ascending max p-value (largest p-value between both species) and the top 5 genes with a mouse log2FC > 1.5 were selected to feature in the dot plot of conserved markers.

**Statistical analysis markers**
Differential gene expression analysis to determine the cluster markers was performed using the Wilcoxon Rank Sum test through the Seurat functions FindAllMarkers and FindMarkers. P-value adjustment was accomplished with Bonferroni correction. RNA markers for the annotated clusters of ChP LV & 4V aggregate were determined with these cutoffs: min.pct = 0.10, min.diff.pct=0.25, logfc.threshold = 0.30, return.thresh = 0.01 and only positive markers were evaluated. For most of the other objects, markers were determined with default cutoffs: min.pct = 0.10, min.diff.pct=-Inf, logfc.threshold = 0.25, return.thresh = 0.01, only.pos = FALSE; except for the Fibroblast origin subset object where the min.pct was set at a strict 0.6 to find highly expressed markers and logfc.threshold was set at 0.30 in the marker analysis on which GO enrichment analysis was run. An extra "score" column was always calculated as a way to rank the importance of the genes as markers. It was calculated with the function: "pct.1/(pct.2+0.01)*avg_logFC". The markers are ordered according to this score.

**Data availability**
All raw sequencing data enclosed in this publication has been deposited in NCBI's Gene Expression Omnibus[12] and will be made accessible after peer-review and publication.

**References**

1. Lun, A. T. L., Bach, K. & Marioni, J. C. Pooling across cells to normalize single-cell RNA sequencing data with many zero counts. Genome Biol. 17, 1–14 (2016).

2. McGinnis, C. S., Murrow, L. M. & Gartner, Z. J. DoubletFinder: Doublet Detection in Single-Cell RNA Sequencing Data Using Artificial Nearest Neighbors. Cell Syst. 8, 329-337.e4 (2019).

3. Korsunsky, I. et al. Fast, sensitive and accurate integration of single-cell data with Harmony. Nat. Methods 2019 1612 16, 1289–1296 (2019).

4. Shah, P. T. et al. Single-Cell Transcriptomics and Fate Mapping of Ependymal Cells Reveals an Absence of Neural Stem Cell Function. Cell 173, 1045-1057.e9 (2018).

5. Vanlandewijck, M. et al. A molecular atlas of cell types and zonation in the brain vasculature. Nature 554, 475–480 (2018).

6. DeSisto, J. et al. Single-Cell Transcriptomic Analyses of the Developing Meninges Reveal Meningeal Fibroblast Diversity and Function. Dev. Cell 54, 43-59.e4 (2020).

7. Zeisel, A. et al. Molecular Architecture of the Mouse Nervous System. Cell 174, 999-1014.e22 (2018).

8. Luecken, M. D. et al. Benchmarking atlas-level data integration in single-cell genomics. Nat. Methods 2021 191 19, 41–50 (2021).

9. Dani, N. et al. A cellular and spatial map of the choroid plexus across brain ventricles and ages. Cell 184, 3056-3074.e21 (2021).

10. Yang, A. C. et al. Dysregulation of brain and choroid plexus cell types in severe COVID-19. Nat. 2021 5957868 595, 565–571 (2021).

11. Polański, K. et al. BBKNN: fast batch alignment of single cell transcriptomes. Bioinformatics 36, 964–965 (2020).

12. Edgar, R., Domrachev, M. & Lash, A. E. Gene Expression Omnibus: NCBI gene expression and hybridization array data repository. Nucleic Acids Res. 30, 207–210 (2002).
